# The Nematode *Oscheius tipulae* as A Genetic Model for Programmed DNA Elimination

**DOI:** 10.1101/2022.08.19.504554

**Authors:** Thomas C. Dockendorff, Brandon Estrem, Jordan Reed, James R. Simmons, Sobhan Bahrami Zadegan, Maxim V. Zagoskin, Vincent Terta, Erin Seaberry, Jianbin Wang

**Author notes:** These authors contributed equally (Thomas C. Dockendorff, Brandon Estrem).

## Abstract

**SUMMARY:** Programmed DNA elimination (PDE) is a notable exception to the paradigm of genome integrity. In metazoa, PDE often occurs during early embryogenesis, coincident with germline to somatic cell differentiation. During PDE, portions of genomic DNA are lost, resulting in reduced somatic genomes. Recent studies have described the sequences lost and chromosome behavior in metazoa. However, a system for studying the mechanisms and consequences of PDE in metazoa is lacking. Here, we established a functional and genetic model for PDE in the free-living nematode *Oscheius tipulae*, a member of the Rhabditidae family, which includes *Caenorhabditis elegans. O. tipulae* was recently suggested to eliminate DNA. Using staged embryos, we show that *O. tipulae* PDE occurs during embryogenesis at the 2-16 cell stages, similar to the human/pig parasitic nematode *Ascaris*. We identified a conserved motif, named <u>S</u>equence <u>F</u>or <u>E</u>limination (SFE), for all 12 break sites on the six chromosomes at the junction of retained and eliminated DNA. SFE mutants exhibit a “fail-to-eliminate” phenotype only at the modified sites. END-seq revealed that breaks can occur at multiple positions within the SFE, with extensive end resection followed by telomere addition to both retained and eliminated ends. We identified through END-seq in the wild-type embryos, genome sequencing of SFE mutants, and comparative genomics of 23 wild isolates a large number of functional SFEs at the chromosome ends. We suggest these alternative SFEs provide flexibility in the sequences eliminated and a fail-safe mechanism for PDE. These studies establish *O. tipulae* as a new, attractive model to study the mechanisms and consequences of PDE in a metazoan.

## INTRODUCTION

Genome integrity is essential to life. Considerable efforts are made to maintain the stability of genomes, as exemplified by elaborate DNA repair mechanisms, cell cycle checkpoints, faithful segregation of chromosomes during mitosis and meiosis, and suppression of mobile genetic elements.^1-4^ Yet genomes also undergo constant changes, often random and small in scale, providing mechanisms for evolution and adaptation. In contrast, programmed DNA elimination (PDE) is a dramatic form of genome change with large amounts of DNA, ranging from 0.5 to 95% of the genome, eliminated during development.^5-9^ PDE is highly selective and reproducible and is an integral part of biology for diverse organisms.

PDE was first discovered in the horse parasite *Parascaris* in the 1880s by Theodor Boveri.^10^ It has since been found in single-cell ciliates,^11,12^ a variety of multicellular organisms across animal phyla,^5,13-16^ and in plants.^17,18^ Initial discoveries of PDE were largely dependent on careful cytological observations, and thus early models for PDE tend to have large genomes and/or discard a large amount of DNA.^5,6^ With the increasing affordability of advanced sequencing technologies, whole genome sequencing has become a sensitive tool to identify PDE.^19^ Genomics has been used to characterize metazoan PDE in depth, including the nature of eliminated sequences^20-27^ and DNA breaks,^20-22^ as well as the evolution of PDE.^15,16,28^ A common theme emerging from these recent genomics studies is the removal of both germline-expressed genes and repetitive sequences during PDE.^5-9^ This suggests that one function of PDE in metazoa is to permanently silence certain germline sequences potentially harmful to somatic cells.^6^ PDE is also used for sex determination,^7,13,15^ remodeling chromosome ends and splitting fused chromosomes,^22^ and appears to be a mechanism for novel gene and chromosome evolution.^6,8,9^

Despite the detailed genomic and cytological descriptions of PDE in many organisms, functional and mechanistic studies of metazoan PDE are limited.^29,30^ Mechanistic insights for PDE, including the contributions and roles of transposable elements and small RNAs, have primarily been gleaned from single-cell ciliates.^11,12,31,32^ The features of PDE in metazoa differ considerably from ciliates. Thus, their underlying mechanisms may differ significantly as well.^6-8^ Unfortunately, most metazoan models for PDE are not well suited for genetic, functional, and/or biochemical analyses. For example, in one of the best-studied organisms, the human and pig parasite *Ascaris*, genetic manipulation is difficult and propagation of the parasite in the pig host is expensive and not sustainable.^33^ While all other known organisms with PDE have unique and interesting features,^5,7,8^ there are also limitations in these systems, including large and complex genomes (copepods, birds, and lampreys), long life cycles (lampreys and birds), and high maintenance (copepods and birds). Genetic and functional tools available for traditional model organisms are not available or very limited for these PDE models.^7,8^ Consequently, concerted genetic, functional, and other molecular analyses have not been carried out on metazoan PDE.

Here, we built upon and extended a previous genomic observation^34^ and established a genetic and functional model for PDE in the free-living nematode *Oscheius tipulae*, a member of the Rhabditidae family, which includes *Caenorhabditis elegans*.^35^ We show that DNA elimination occurs during early embryogenesis in *O. tipulae*. Our comprehensive RNA-seq analyses reveal expression profiles of the eliminated genes and differentially expressed genes during DNA elimination. Notably, we identified and characterized a conserved sequence motif associated with the DNA break sites and demonstrated its direct role in DNA elimination. DNA breaks occur within the motif, followed by end resection and telomere healing. Additional breaks occur simultaneously in the eliminated DNA, perhaps serving as a fail-safe mechanism for PDE. We revealed the abundance and variations of this motif in 23 *O. tipulae* wild isolates from around the world. The ability to utilize the available and developing tools of genomics, biochemistry, molecular biology, cell biology, and genetics, along with its small genome and short life cycle, make *O. tipulae* an excellent model for studying the molecular mechanisms of PDE in a multicellular organism.

## RESULTS

### *O. tipulae* undergoes PDE during early embryogenesis

*O. tipulae* is best known as a comparative model to *C. elegans* for studies on the evolution of developmental pathways.^35-38^ *O. tipulae* was suggested to eliminate DNA in a recent genomic study from The Tree of Life project at the Sanger Institute,^39^ where Gonzalez de la Rosa et. al. assembled a telomere-to-telomere genome of *O. tipulae* and found a lower coverage of sequencing reads at the ends of all six chromosomes.^34^ This data was reminiscent of DNA elimination in the parasitic nematode *Ascaris*, where the ends of all 24 germline chromosomes are removed and remodeled.^22^ The decreased read coverage at chromosome ends (compared to the complete loss of reads), was consistent with the analysis of a mixed population of worms (embryos, larvae, and gravid adults) used in the sequencing experiments, thus reflecting the presence of both germline and somatic cells in the samples.^19,20^

To confirm that PDE indeed occurs in *O. tipulae*, we collected (see Methods and Fig. S1) and sequenced genomes from the germline (mainly 2-4 cell embryos), multiple stages of early embryogenesis, and somatic cells (L1 larvae) (Fig. 1). We found that in the 2-4 cell sample, the reads at all ends of chromosomes have a higher genome coverage than in later stages of development. This coverage (65%, compared to retained regions of the genome) is consistent with PDE starting at the 2-cell stage with some potential residual DNA from the eliminated regions. The coverage decreases progressively through embryogenesis to ∼4% in the L1 stages (Fig. 1 and Fig. S2A). These changes in genome coverage are proportional to the percentage of germ cells in the sample. We could not obtain 100% or 0% read coverage due to the somatic contamination in the 2-4 cell sample (Fig. S1 and Fig. S2A) and the presence of two primordial germ cells (PGCs) and underdeveloped embryos in the L1 sample. Likewise, a mixed worm population containing largely somatic cells consistently showed only ∼10% sequencing coverage of the eliminated ends of the chromosomes. The progressive drop of read coverage indicates the loss of DNA in the somatic cells, suggesting PDE occurs during early embryogenesis (Fig. 1).

**Figure 1.**
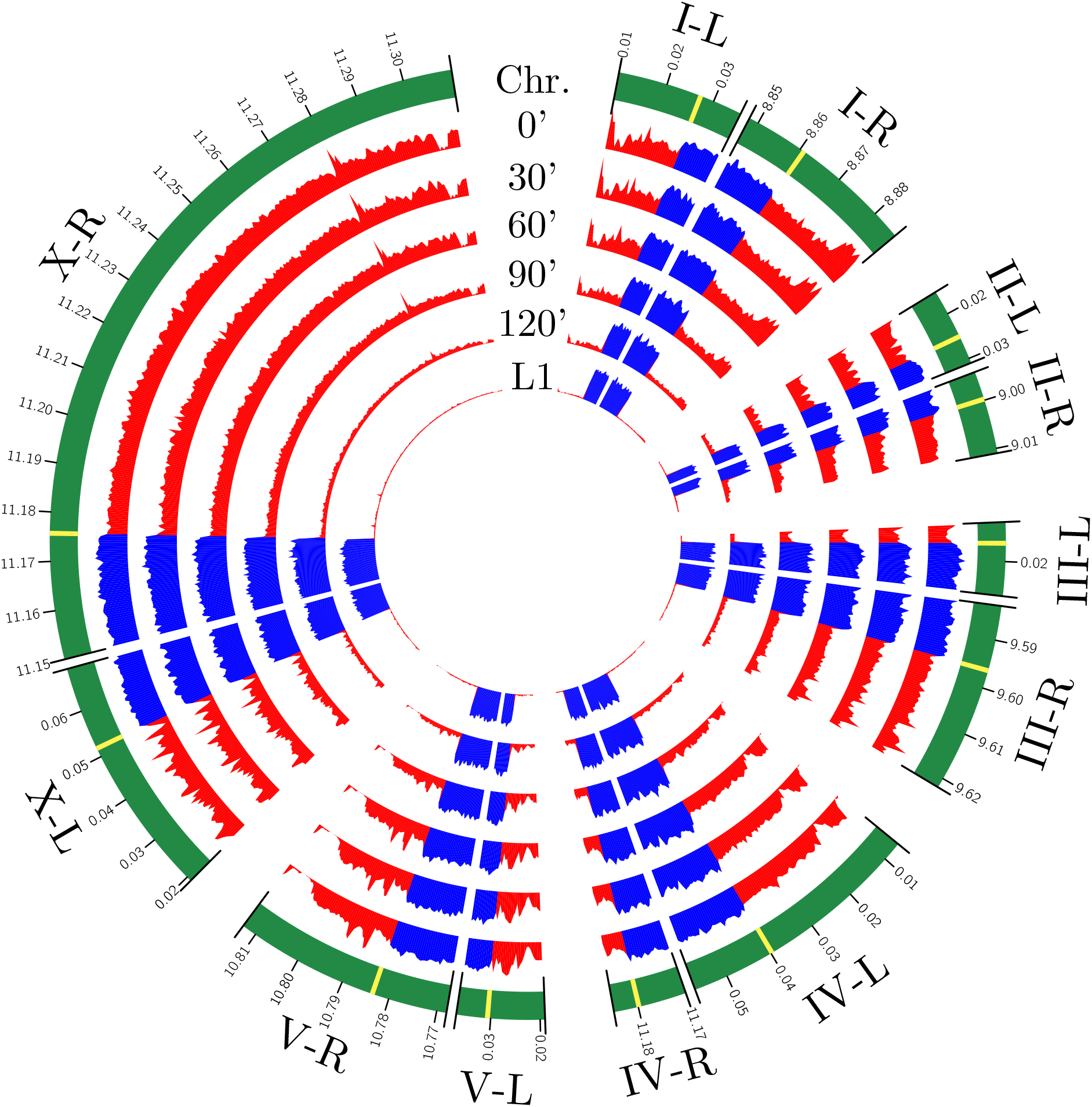
*O. tipulae* undergoes PDE during early embryogenesis. Genomic DNA at the ends of *O. tipulae* chromosomes is lost during early development. A Circos^96^ plot shows the Illumina read coverage for early embryos (time indicates minutes of development after egg harvest, 0’ = 2-4 cells, see Figure S1) and L1 larvae. The retained DNA is blue and the eliminated DNA is red. For simplicity, the majority of retained DNA is not shown (indicated as a gap, ||, in the middle of each chromosome). The junctions of retained and eliminated DNA are colored in yellow (outer circle). Genome coverage is plotted in 500-bp bins, and the coordinates are displayed in Mb.

In all studied ascarid nematodes, PDE also occurs during early embryogenesis. However, the exact timing of PDE varies.^40^ For example, in *Parascaris*, PDE occurs during the 2-32 cell stages, while in *Ascaris*, it occurs in 4-16 cell embryos.^13^ To determine the timing of PDE in *O. tipulae*, we used the cell numbers in the staged embryos to determine the developmental timing of PDE that best fits our sequencing coverage (Fig. S2B). This analysis revealed that PDE in *O. tipulae* most likely occurs during the 2-16 cell stages. The timing of PDE is further supported by Hoechst staining, which detects the presumptive eliminated DNA foci in the cytoplasm as early as the 4-cell stage (Fig. S3), suggesting PDE occurs prior to 4-cell embryos. Thus, our sequencing and staining data demonstrate that *O. tipulae* undergoes PDE during early embryogenesis at the 2-16 cell stages.

### Expression of *O. tipulae* eliminated genes

Previous work identified 115 genes in the eliminated regions, including many helitron-like genes.^34^ However, the gene models were primarily based on *in silico* prediction, and expression data for *O. tipulae* genes was lacking. To determine the expression of retained and eliminated genes, we built comprehensive transcriptomes for *O. tipulae* by carrying out RNA-seq on 20 developmental *O. tipulae* samples (Fig. 2A and Table S2). These samples include early embryogenesis with staged embryos during and after PDE, as well as samples from gastrulation, morphogenesis, staged larvae (L1 – L4), dauer, post-dauer larvae, young adults, and mature adults (Fig. 2A). Studies in ascarid nematodes indicated that many of the eliminated genes were expressed during spermatogenesis.^20-22^ Thus, we also sequenced RNA from male worms (handpicked from a high incidence of males [HIM] mutant) to identify male-enriched genes (Table S3 and S4). Using the RNA-seq data, we refined the gene models for *O. tipulae*. Major improvements in the gene models include the comprehensive extension of 5’ and 3’ UTRs, identification of alternatively spliced isoforms, 560 new genes, and a reduction of 1,809 genes following the merging/collapsing of gene models (Fig. S4). Overall, this RNA-based gene prediction identified 14,558 genes, 36,790 transcripts, and 112 eliminated genes (see Table S3).

**Figure 2.**
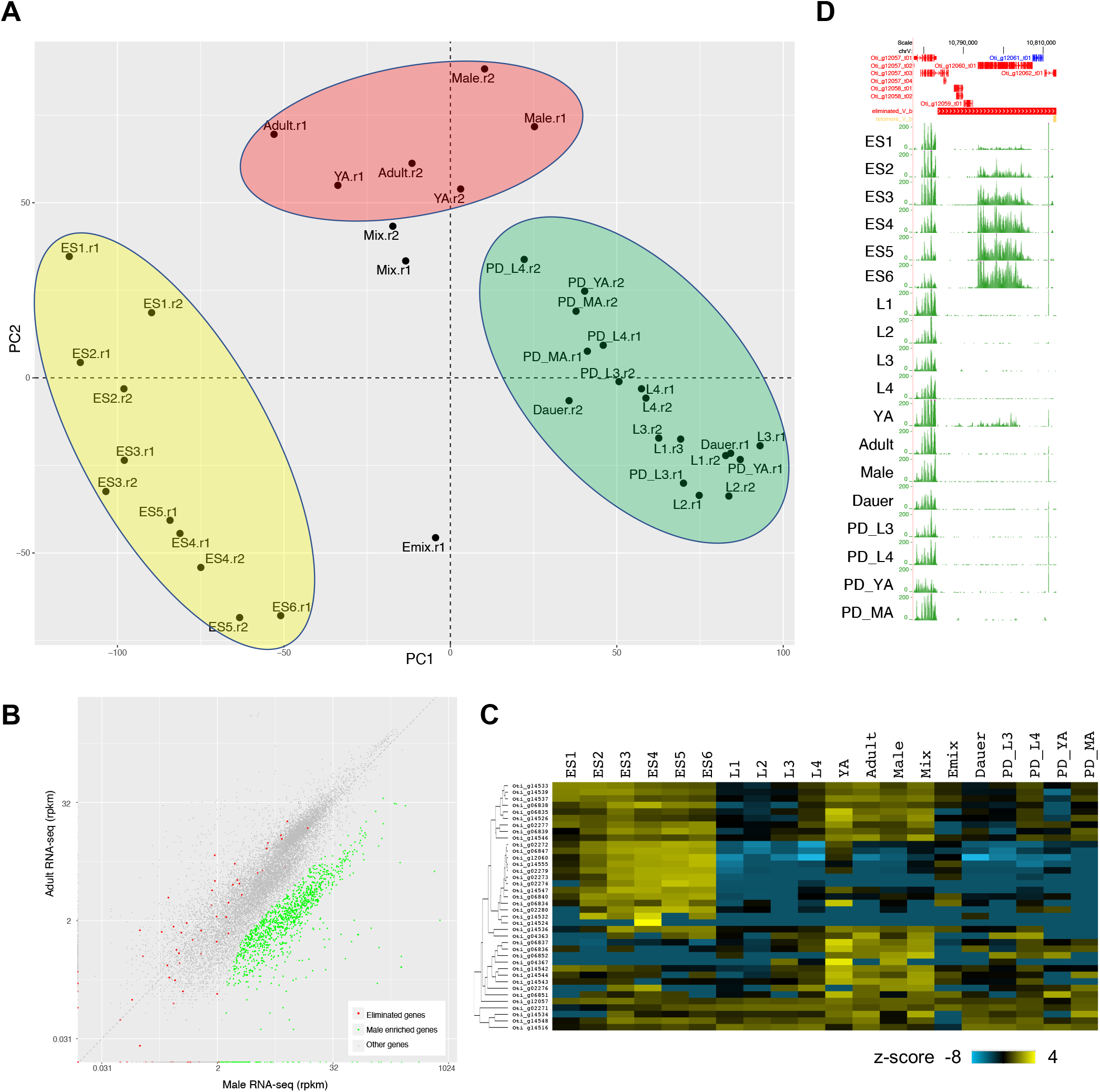
*O. tipulae* eliminated DNA encodes germline- and early embryo-expressed genes. **A**. Comprehensive RNA-seq for *O. tipulae* embryos and other developmental stages. The Principal Component Analysis plot shows the relationship of the RNA expression profiles among the sampled developmental stages and replicates (labeled as r1 and r2). Sample descriptions: ES indicates embryo stage, with ES1 = 2-4 cells, ES2 = 4-8 cells, ES3 = 8-16 cells, ES4 = 16-32 cells, ES5 = 32-64 cells, ES6 = 64-128 cells, and ES7 = 128-256 cells; PD = post-dauer; YA = young adult; MA = mature adult; Emix = mix of embryos; and Mix = mix of embryos, larvae, and adults (see Fig. S1 and Table S2). Note the clustering of the embryos (yellow), larvae and post-dauer worms (green), and mature worms (red). **B**. Eliminated genes are not enriched in the males. Comparison of RNA expression (rpkm) between males and hermaphrodites revealed 1,229 male-enriched genes (blue). None of them overlap with eliminated genes (red). **C**. Heatmap showing the expression profiles of 36 eliminated genes. The average rpkm values from the replicates across the stages were converted to z-score. The other 74 eliminated genes have little to no expression across these developmental stages (rpkm < 2). **D**. Genome browser view of an eliminated gene from chrV-R, showing elevated expression during embryogenesis.

One-third (38) of the eliminated genes are expressed in the germline or early embryos (Fig. 2C and Table S3). However, the expression level of these genes is on average lower than retained genes (Table S3). This could be due to the reduced level of germline sequences in these mixed samples and/or that most of the eliminated genes are not transcriptionally active as they may be close to heterochromatic regions at the ends of the chromosomes. Consistently, the other two-thirds (74) of the eliminated genes have little or no expression in all sampled stages (maximal rpkm < 2). About half of the eliminated genes are testis-specific in ascarids.^20,21,41^ We used RNA-seq data from the male worms to determine if *O. tipulae* also eliminates male-specific genes. We identified 1,229 male-enriched genes (5x male over mature hermaphrodite, p-value < 0.05; Fig. S5 and Table S4), including major sperm proteins and the Argonaute ALG-4.^42,43^ However, none of the 112 eliminated genes is enriched in the male sample (Fig. 2B). To investigate the potential function of these eliminated genes, we annotated the genes and performed gene ontology (GO) and tissue enrichment analysis.^44,45^ The tissue distribution analysis shows that expression of the eliminated gene is enriched in *C. elegans* Z2 and Z3 germ cells. However, no GO enrichment was identified due to the low number of eliminated genes and the presence of many hypothetical genes (Table S3). This is consistent with the observations in other nematodes where many eliminated genes are “novel” genes.^6,41^

Surprisingly, eight genes from the eliminated regions are highly expressed in the 64-128 cell stage (Fig. 2D). This is a stage where all PDE events have been completed, as is evident by our genome sequencing data (Fig. 1). These genes do not have extra copies in the retained regions of the genome. Since these RNAs are not highly expressed in earlier stages, they must be newly transcribed – likely from the two PGCs. Considering that the two PGCs are a small fraction of the total cells at this stage, the expression level for these genes in the PGCs may be ∼50x higher than the reported values. Gene annotation indicates that these genes are orthologous to the ATP-dependent DNA helicase (*pif-1*) in *C. elegans*. These eight genes contain a helitron-like domain and have five copies in the eliminated regions of the *O. tipulae* genome. Altogether, our RNA-seq data and functional annotations suggest that the eliminated genes are expressed in the germline or early embryos and are not expressed in males. A few genes have a burst of transcription in the primordial germ cells. In addition, many eliminated genes have no predicted functions, consistent with observations in other nematode systems,^6,46^ suggesting they may have evolved recently.

### Differentially expressed genes during *O. tipulae* PDE

Using RNA-seq data, we defined differentially expressed genes during early embryogenesis where PDE occurs (Fig. S5 and Table S4). We predict that genes involved in PDE will be highly expressed in the 2-16 cell embryos but down-regulated in the 16-128 cell embryos. Using comparisons between 2-8 cell vs. 8-32 cell and 2-16 cell vs. 16-128 cell, we identified 622 genes that are consistently enriched (2x fold change and p-value < 0.05 in both comparisons) during PDE stages (Fig. S5 and Table S4). GO analysis suggests that these genes are enriched in meiotic cell cycle function, macromolecule catabolic processing, reproduction, pole plasm, and ribonucleoprotein granules (Table S4). In comparison, 1,317 genes are upregulated in late embryos compared to early embryos; they are enriched in neuron development, regulation of cell motility, cell morphogenesis, and cell projection organization (Table S4). Because early embryogenesis requires complex and orchestrated regulation of gene expression, we reason that many enriched genes are involved in early developmental processes. However, we also expect that genes involved in PDE are enriched in the early embryos. For example, the expression of the telomerase gene (Oti_g07275) is upregulated during early embryogenesis and peaks at the 8-16 cell stage, consistent with its role in telomere healing during PDE. This comprehensive developmental RNA-seq expression profile provides a critical genomic resource for future genetic, molecular, and developmental studies of PDE and in *O. tipulae* biology in general.

### A conserved motif at *O. tipulae* break regions

PDE reproducibly leads to DNA breaks at specific regions of nematode chromosomes. How does the cell determine where the DNA breaks should occur? In *Ascaris*, telomere healing, and likely the DNA break, occurs with high fidelity and heterogeneously within 3-6 kb regions called chromosomal breakage regions (CBRs).^20-22^ Sequence analysis has identified no conserved motifs within these CBRs.^21^ In contrast, examination of *O. tipulae* sequences at the junction of the retained and eliminated DNA reveals a highly conserved motif (Fig. 3A-B). This motif, named <u>S</u>equence <u>F</u>or <u>E</u>limination (SFE, or pronounced as SAFE), spans about 30 bp and is present at the 12 sites on both ends of the six chromosomes. Since these 12 sites determine where the DNA breaks occur and what distal sequences are eliminated in the wild type CEW1 strain, we define these as canonical break sites (see below for alternative sites used at the ends of the chromosomes). The most conserved bases of the motif are arranged in a palindrome (Fig. 3B) – a common recognition signal for a variety of DNA-binding proteins, including those that cleave or modify DNA. The strong conservation of this motif at the DNA break sites suggests that the SFE motif may participate in the DNA break and/or telomere addition processes.

**Figure 3.**
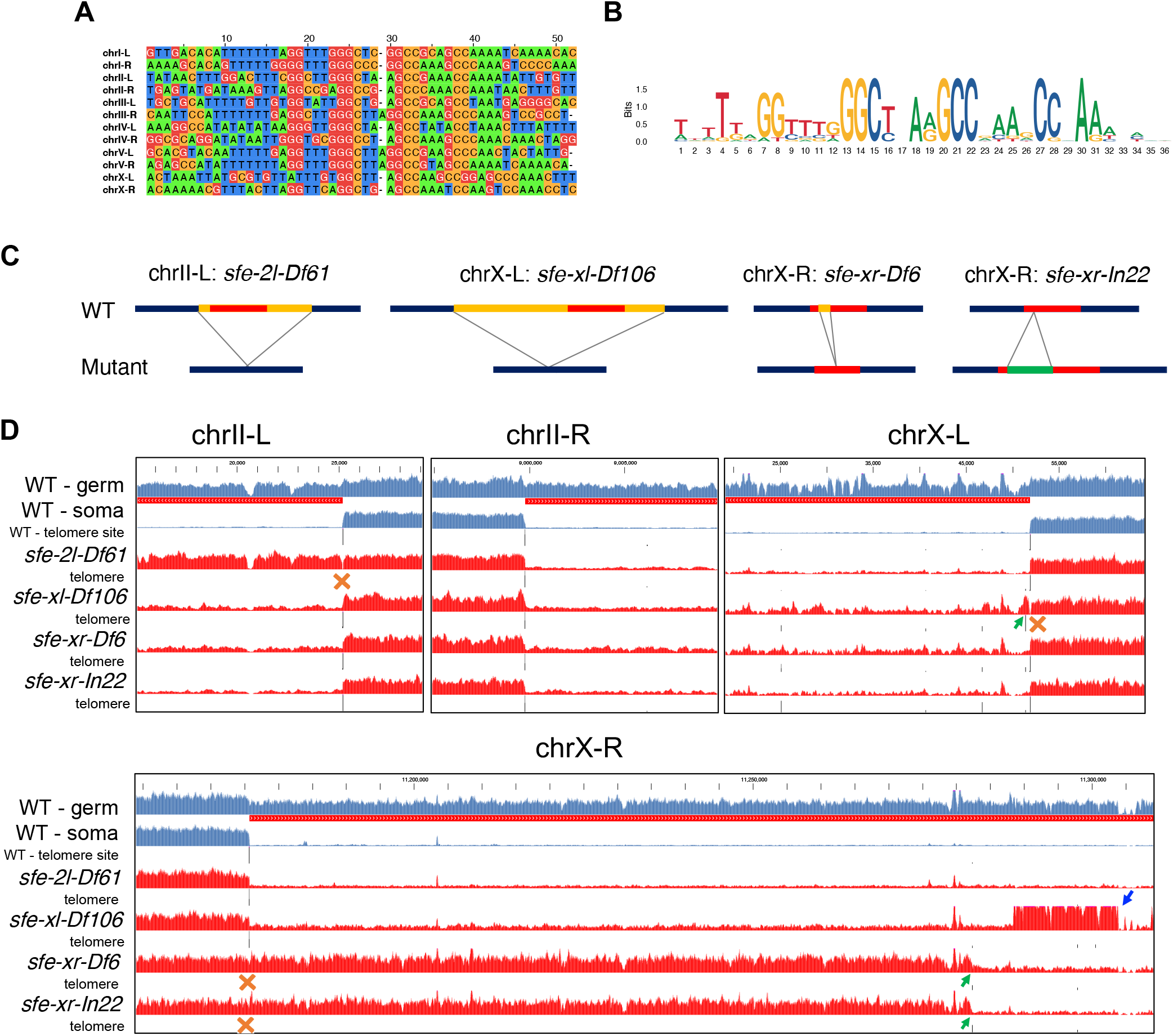
Conserved motif at *O. tipulae* break region is essential for DNA elimination. **A**. Alignment of 12 sequences from the junctions of the retained and eliminated DNA (canonical SFE sites; these sites define the somatic telomere ends in the wild type CEW1 strain) using Jalview^97^ with manual adjustment. **B**. Motif for the canonical SFEs using ggseqlogo.^98^ **C**. Schematic design of the genome editing for the SFEs. Yellow = deleted DNA, red = core motif region, and green = inserted DNA. **D**. “Fail-to-eliminate” phenotype confirmed by genome sequencing. Four ends of the *O. tipulae* chromosomes are shown (labeled at the top). For each end, genome coverage from the wild-type germline and somatic cells (blue), as well as sequencing reads from all four mutants (red), was shown. The eliminated regions are marked as red bars below the germline track. Also shown are the telomere addition sites derived from the sequencing data. The mutated sites are indicated with orange Xs. At each mutated site, either a complete (chrII-L) or a partial (chrX-L and chrX-R) “fail-to-eliminate” phenotype was observed. Green arrows point to the alternative sites used in the mutants. The blue arrow indicates DNA with an elevated read coverage that is supposed to be eliminated in the somatic cells. chrII-R serves as a control site where no SFE has been modified. The genome coverage for the eliminated regions varies due to the variations of germ cells percentage in the sequenced samples.

### Mutations in the SFE motif result in a fail-to-eliminate phenotype

To explore the role of the SFE motif in the DNA elimination/telomere addition process, we used CRISPR-Cas9 to remove or alter the motif at three break sites. These sites were chosen due to their variations in the amounts of non-telomeric sequences lost (chrII Left = 10.1 kb, chrX Left = 32.7 kb, and chrX Right = 133.5 kb). Using a “roller” phenotype as a co-CRISPR marker (see Materials and Methods and Fig. S10), we obtained four SFE mutants with disrupted motifs (Fig. 3C). These mutants were confirmed by PCR analyses and Sanger sequencing (Fig. S6). All four SFE homozygous mutants are viable and fertile. However, this does not preclude the possibility of subtle phenotypes in these single mutants or that mutations to additional SFEs might result in more apparent phenotypes. Illumina sequencing of genomic DNA from mixed populations (eggs, larvae, adults) shows that while mutations of this consensus motif block elimination at the altered sites, alternative elimination sites within the eliminated DNA were often used. These sites, closer to the ends of the chromosomes, led to elimination of smaller regions of the chromosome end (Fig. 3D). For example, two independent mutations within the SFE motif at the right end of chrX, *sfe-xr-Df6* and *sfe-xr-In22*, resulted in the retention of 106.6 kb of the typically eliminated 133.5 kb of DNA (Fig. 3D, alternative SFE indicated by green arrows in chrX-R). Furthermore, a mutation that removes the SFE motif from the left end of chrX (*sfe-xl-Df106*) prevented the elimination of 473 bp due to an alternative SFE (Fig. 3D, green arrow in chrX-L). The use of these alternative sites suggests that although there may be preferred sites, the alternative sites may serve as a fail-safe mechanism to ensure the removal and remodeling of the chromosome end (see below END-seq results). Overall, these results show that the SFE is a key element involved in the elimination process, providing a sequence for the DNA break and telomere addition. We note, however, that not all altered SFE sites lead to the use of an alternative break site. The SFE mutant on the left end of chrII (*sfe-2l-Df61*) resulted in the retention of the entire small 10.1 kb subtelomeric region and its telomeric DNA.

### *O. tipulae* DNA break ends are resected and healed with telomere addition

To further characterize the DNA breaks in *O. tipulae* PDE, we adapted an END-seq approach that identifies DNA double-strand breaks (DSBs) and the resection of DSB ends.^47,48^ We demonstrated that END-seq can identify exogenously introduced FseI or AsiSI DSBs with resected ends (3’-overhangs) at single nucleotide resolution in *O. tipulae* early embryos (Fig. 4A-B). Our END-seq revealed that PDE breaks occur within the conserved SFE motif, followed by extensive end resections (an average of 1.4 kb) (Fig. 4C). Within the motif, we observed overlapping END-seq signals between the retained and eliminated ends of the breaks, suggesting that the DNA breaks can occur at various positions within the motif. END-seq can also capture new telomere addition events (as long as the telomere length does not exceed the length of Illumina reads; see Methods). Our analysis revealed that telomere addition occurs on both the retained and eliminated ends with equal read frequency (Fig. 4C-D), suggesting an initial unbiased telomere healing process to both ends of a break. Strikingly, >98% of the telomere addition sites are within the GGC/GCC position in the center of the palindromic motif (Fig. S7). We conclude that this GGC/GCC sequence serves as the primer for synthesizing new telomeres (**<u>GGC</u>**TTA/TAA**<u>GCC</u>**)_n_ after the break. The other 2% of the telomere addition sites are within the motif, but often have only two nucleotides that match the telomere sequence, presumably providing alternative primer sites for telomerase (Fig. S7).

**Figure 4.**
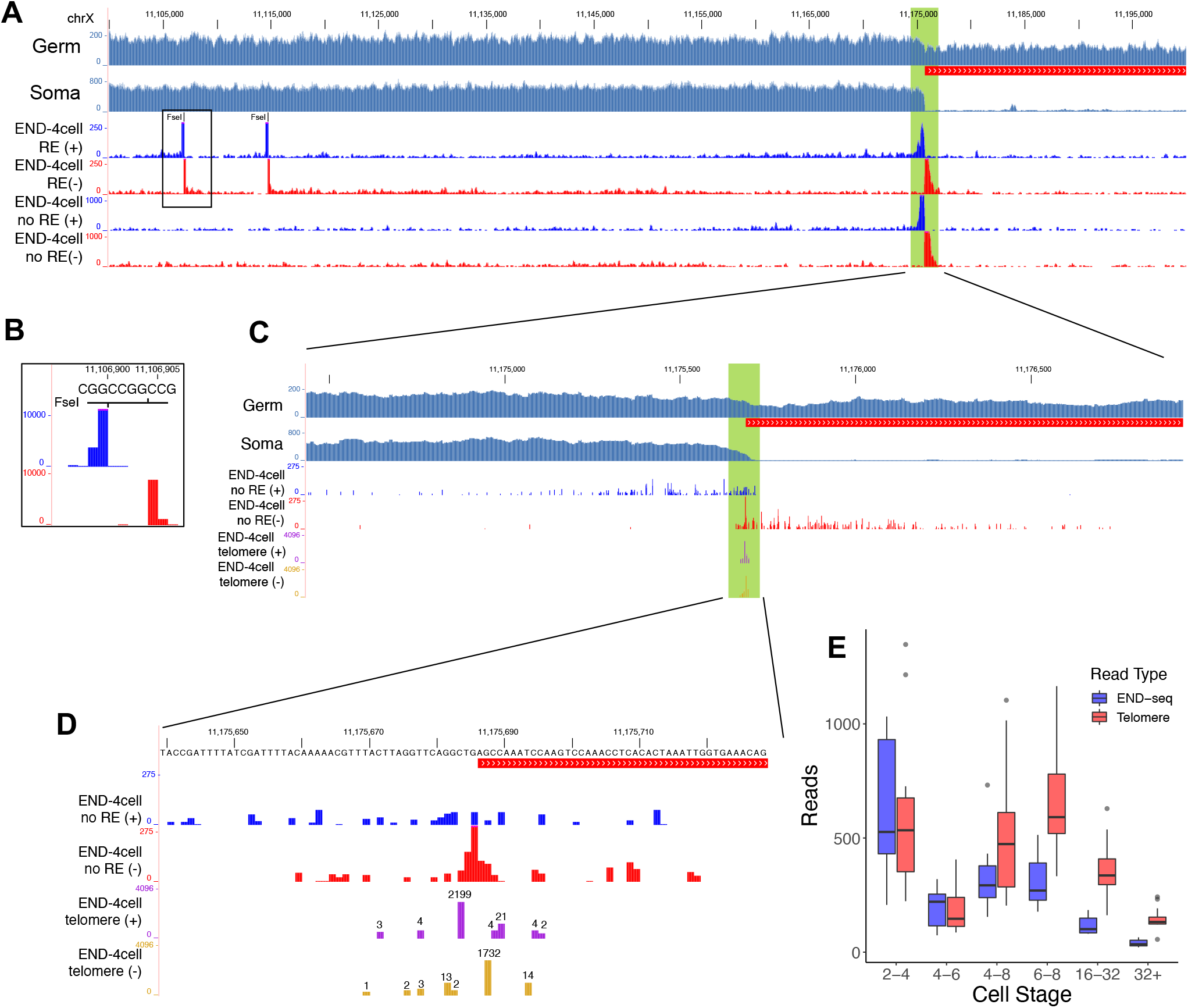
*O. tipulae* break ends are resected and healed with telomere added at both ends. **A-D**. END-seq data from a population of 2-4-cell embryos showing features of the DSBs. **A**. Genome browser view of full END-seq reads from a of 100-kb region at the right end of chrX containing two FseI restriction sites and an SFE site (chrX-R). Germline and somatic read coverage are shown to indicate the break site and the eliminated DNA (red line). END-seq libraries with or without FseI restriction digestion are labeled as RE or no RE. The reads are separated into the forward (+) and reverse strand (-). The END-seq read coverage is shown. Highlighted green region (2.5 kb) at the break site is zoomed in **C. B**. Inset of one FseI site. Libraries treated with FseI show high enrichment of END-seq reads at the FseI sites. The gap between the two strand is caused by the 4-nt 3’-overhang that is removed during END-seq blunting procedure (see Methods). The coverage for the end of the reads is shown. **C**. Zoomed in view of the 5’-end of END-seq reads at the SFE. Counts for the end of the reads accumulate in the tails on both sides of the break, consistent with the end resection pattern at a DSB.^47,48^ The END-seq signal is more enriched at the core motif region. The overlapping signals from the two strands cannot be derived from a single break site (see Fig. 4B), indicating some DSBs can occur at other sites within the overlapping region. Also shown are the telomere addition sites (purple and gold tracks) enriched in the END-seq data (see Methods). **D**. Zoomed in view of a 90-bp region containing the core motif, showing the sequence, 5’-ends of END-seq reads, and the frequency of telomere-unique reads (in log 2 scale, with the number on the top of the site). Note the close-to-equal frequency of telomere addition at the both the retained and eliminated ends, with the most common telomere addition sites at GGC/GCC. **E**. Delay of telomere addition after the DSBs. The number of reads for END-seq breaks (blue) and telomere addition (red) within 1 kb on either side of the SFE are shown for the early developmental stages. Note that the timing of telomere addition shows a delay relative to DSB formation.

The existence of multiple telomere addition sites also explains the overlapping END-seq signals that suggest DNA breaks can occur at multiple sites within the SFE motif (Fig. 4C). In addition, END-seq data through early developmental stages shows that the 2-4 cell stage has the highest number of breaks, with the frequency of breaks greatly reduced at the 16-32 cell stage (Fig. 4E and Fig. S8). These data are consistent with the timing of *O. tipulae* PDE at the 2-16 cell stage (Fig. 1). Notably, there is a temporal lag between when the DNA breaks occur and the addition of the telomeres (Fig. 4E and Fig. S8), suggesting a regulated transition between DNA break and telomere healing.

### Additional break sites in the eliminated regions provide a fail-safe mechanism for PDE

Surprisingly, in addition to the 12 canonical SFE sites at the junctions of retained and eliminated DNA in the wild-type CEW1 strain, analysis of our END-seq data also identified 12 alternative sites in the eliminated regions (Fig. 5). These break sites would be largely missed by the genome sequencing of somatic tissues^19^ as they are transient in nature and lost together with the entire eliminated regions in the somatic cells. END-seq enriches these signals by capturing the ends of DNA breaks during PDE as these breaks always occur. The read frequency, end resection pattern, and features of telomere addition for these 12 alternative SFE sites are the same as the 12 canonical SFE sites (Fig. 5A-B). Although the sequences for these alternative SFEs exhibit slightly variations, they all share the conserved motif (Fig. 5C-D). Interestingly, two groups of alternative SFEs, one with six sites and the other with three, share high sequence similarity within each group, suggesting they were recently duplicated (Fig. 5B-C). The distribution of these alternative sites is biased towards the ends of the chromosomes, although not all chromosome ends have an alternative site (Fig. 5B). Our END-seq data and analyses suggest that during the onset of PDE, all 24 breaks occur simultaneously in the *O. tipulae* genome.

**Figure 5.**
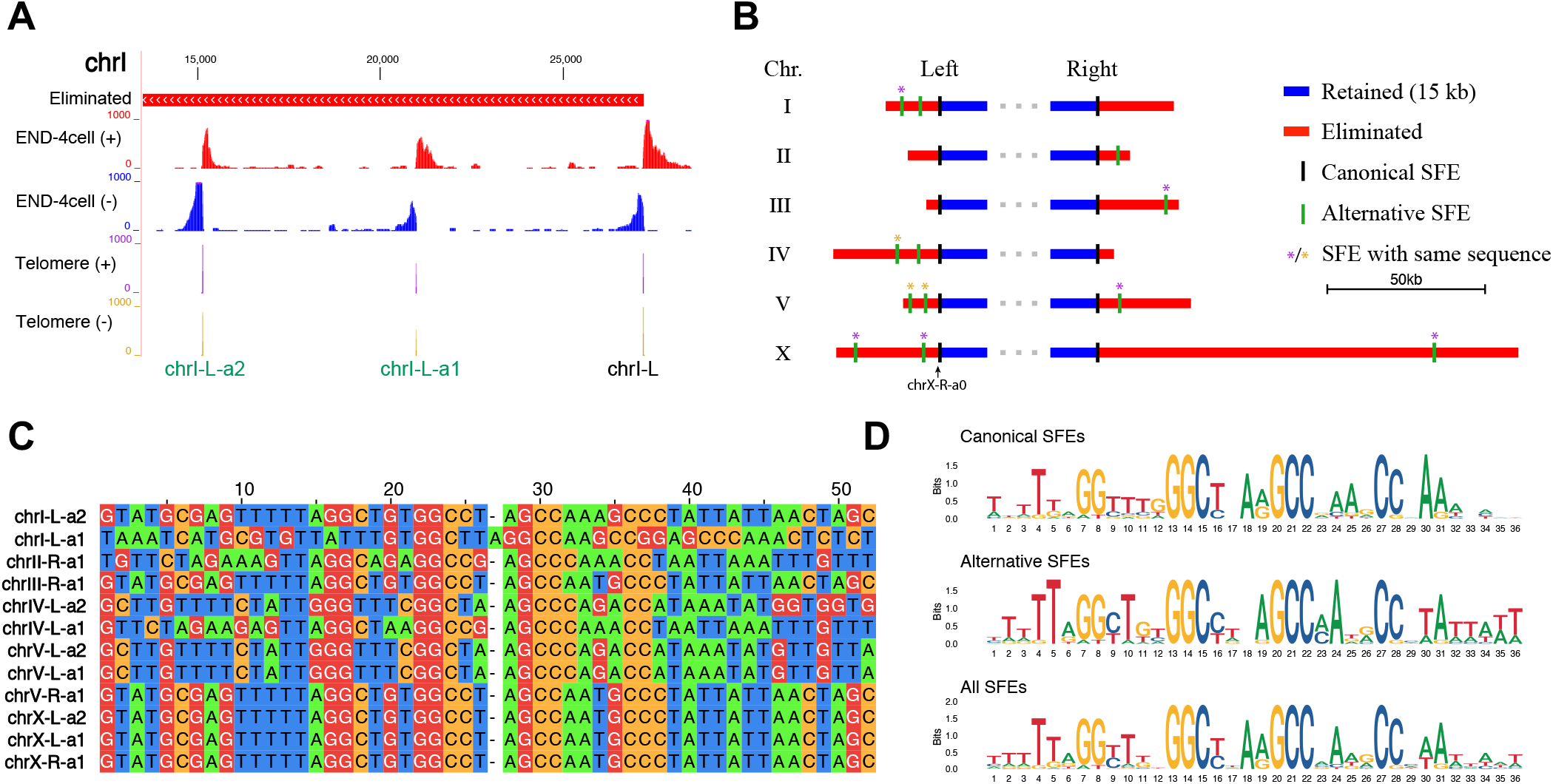
Twelve additional sites in the eliminated regions provide a fail-safe mechanism. **A**. END-seq reveals additional break sites in the eliminated regions. Shown is the left end of chrI, where two additional SFEs (chrI-L-a1 and chrI-L-a2) are identified. They have similar END-seq resection patterns and telomere addition reads compared to the canonical SFE (chrI-L). **B**. The distribution of the 12 alternative SFEs at the ends of the chromosomes. Most of the retained DNA is not shown (represented by dots). Two groups of SFEs with similar sequences are marked with * in purple or gold. These are *bona fide* sites as their read frequencies are at the same level (not lower) as the canonical sites. **C**. Alignment of 12 sequences from the alternative SFE sites. **D**. Motifs derived from canonical and/or alternative SFEs. Note the presence of the conserved GGC/GCC sites for the palindromic sequences.

These alternative SFEs may act as a fail-safe mechanism for *O. tipulae* PDE to ensure that the chromosome ends are remodeled. Indeed, in our SFE mutants at chrX-R (*sfe-xr-Df6* and *sfe-xr-In22*), one of the alternative SFE is used when the original SFE is mutated (Fig. 3D and Fig. 5B). Interestingly, in the SFE mutant at chrX-L (*sfe-xl-Df106*), a site not identified in our END-seq data, was used to make an alternative break (Fig. 3D and 5B). In sum, our data suggest that additional DNA break sites exist to provide a fail-safe mechanism for PDE. Future studies are needed to identify factors that determine the choice of SFEs in different scenarios, including various mutants and wild isolates (see below).

### *O. tipulae* break sites are not more chromatin accessible during PDE

In *Ascaris*, all chromosomal breakage regions become more chromatin accessible during and after PDE.^21,22^ We wondered if *O. tipulae* has similar epigenetic features associated with the SFE sites. We used ATAC-seq to determine the chromatin accessibility during *O. tipulae* early embryogenesis (Fig. S9). While ATAC-seq provided a rich trove of information, we did not find an association between open chromatin and the SFEs. The ends of *O. tipulae* chromosomes are in general not more chromatin accessible than the middle of the chromosomes. For some sites, the chromatin becomes slightly more accessible after PDE, but this open chromatin is not found across all sites (Fig. S9). This difference in accessibility at the break sites suggests that different mechanisms may be involved for break site recognition and/or DNA breaks in the two species. In *O. tipulae*, the break sites (SFEs) are specifically targeted, while in *Ascaris*, the open chromatin may allow the breaks to occur within a broad region of the CBRs (see Discussion).

### *O. tipulae* break sites are flexible in wild isolates

The breaks and alternative breaks described above are from the *O. tipulae* CEW1 strain and its mutants. To assess the conservation and divergence of these break sites in other natural strains, we obtained 23 wild isolates of *O. tipulae* (a gift from Marie-Anne Félix, IBENS; Table S5) from geographically diverse locations around the world (Fig. 6A). For each isolate, we used Illumina to sequence the genomic DNA from a mixed population (eggs, larvae, and adults) that contains ∼5-10% of germline cells. *De novo* assemblies of the complete mitochondrial genomes were used to build a phylogenetic tree (Fig. 6B). This tree is consistent with the geographical distribution, establishing the evolutionary relationship among the strains.

**Figure 6.**
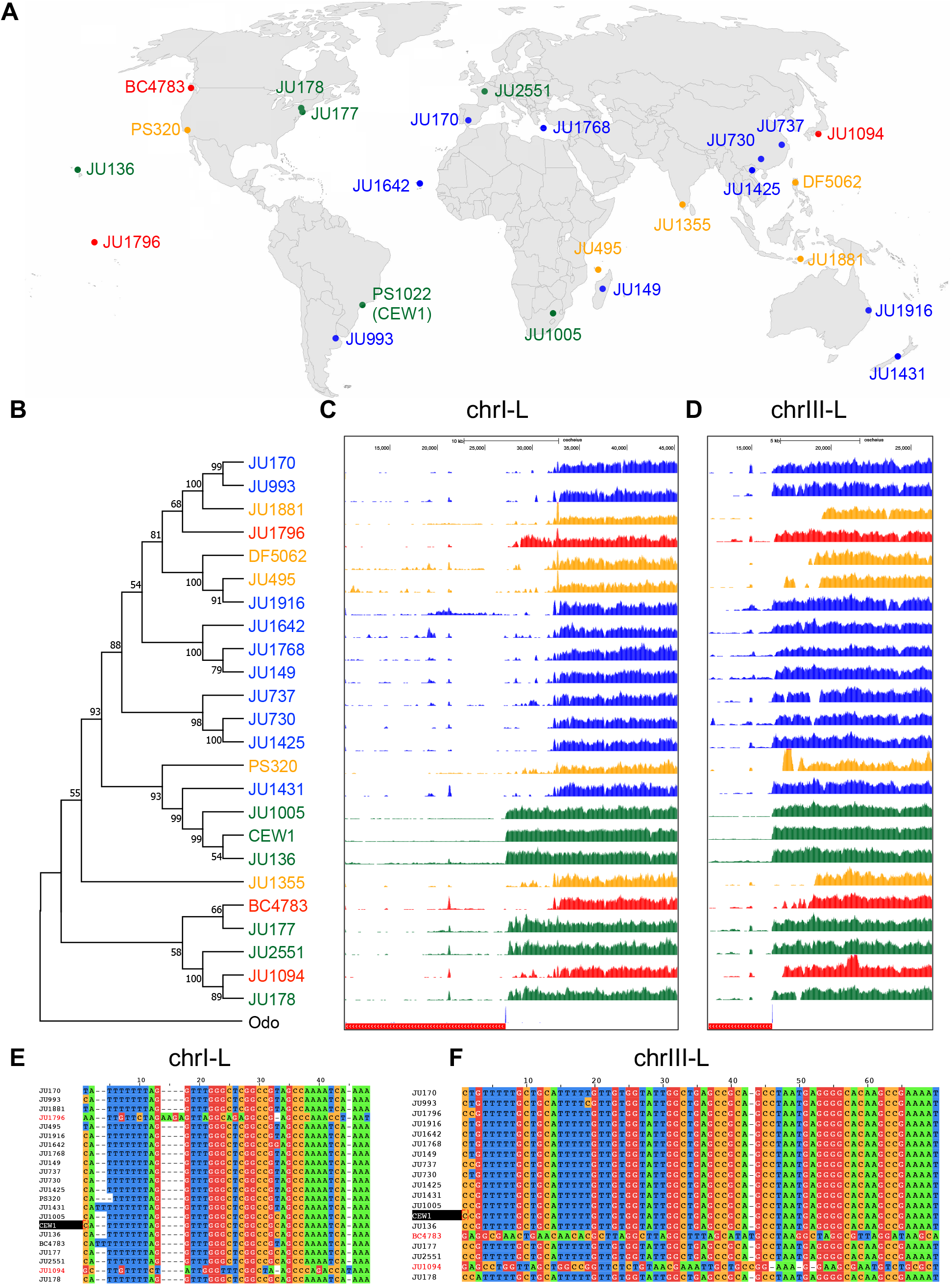
*O. tipulae* break sites are flexible in wild isolates. **A**. Wild isolates of *O. tipulae* from around the world used in this study (strains were from M. A. Félix; see Table S5 for descriptions of the strains). The strains were colored based on their phylogenetic relationships and sequence features associated with PDE (see **B-F**). **B**. A phylogenetic tree based the whole mitochondria genomes. The tree was constructed using MEGA11^95^. The mitochondrial genome of *Oscheius dolichura* (Odo, accession #: OW051498.1) was used as an outgroup for the tree. **C** and **D**. Variations of the potential break sites and eliminated DNA across the strains. Genomic reads from all strains (including CEW1) were mapped to the CEW1 reference genome. The telomere addition sites (blue) and eliminated DNA (red) for CEW1 are indicated at the bottom. Eliminated DNA in CEW1 genome is marked at the bottom red bar (left sides), with telomere addition sites at the junctions (in blue). **E** and **F**. Alignment of *de novo* assembled sequences at the break sites for chrI-L (**E**) and chrIII-L (**F**). CEW1 is highlighted in black. Strains with notably different sequences to CEW1 are colored in red. Note that not all strains have assembled or identified sequences at these break sites.

We mapped the genomic reads from these wild isolates to the CEW1 reference genome to examine the conservation and variation in chromosome end remodeling on the six chromosomes. Two break sites (chrI-L and chrIII-L) show extensive variations among the wild isolates (Fig. 6C-D). These variations would result in the elimination of more sequences (1-5 kb) in these wild isolates compared to the CEW1 strain. However, this mapping-based analysis might be biased if the genomes of these strains differ substantially from the reference CEW1. To reduce this potential bias, we used *de novo* assemblies to further evaluate the differences observed (see Methods). We assembled telomere addition sites for each strain at chrI-L and chrIII-L to identify potential SFEs. For the 21 wild isolates with assembled sequences at chrI-L, two (JU1796 and JU1094) have notably different sequences at chrI-L compared to the SFEs from other strains (Fig. 6E). Similarly, two (BC4783 and JU1094) out of the 18 assembled strains for chrIII-L have different SFE sequences (Fig. 6F). While these sequences have deviated from the CEW1 reference at the two sites, they all have the core conserved SFE motif, reinforcing the idea that the motif is essential for the break and telomere addition.

### Widespread distribution of potential break sites at the ends of CEW1 chromosomes

We next determined if the divergent SFEs (two for chrI-L and two for chrIII-L, Fig. 6E-F) from the wild isolates were present in the CEW1 genome. Surprisingly, all four sequences can be mapped to the CEW1 genome, but are in different regions of the CEW1 genome than determined for the wild isolates. This suggests that these sites may have been rearranged (reshuffled) during the evolution of the genomes. This also indicates that the CEW1 genome potentially harbors SFEs that are not used during PDE. Using genomic reads from CEW1 and the other 23 strains, we identified 399 potential SFEs in the CEW1 genome (see Methods; Table S6). Of them, 12 are the canonical breaks, 66 are sites present in multiple strains, 76 are sites unique to a single strain but with high read frequency (> 5/10 million reads), and the remaining sites (245) are unique to a single strain and have low read frequency (see examples in Fig. 7A). Because these 399 SFEs are from only 24 strains, we predict that the number of potential SFEs will be higher if more strains are examined. Indeed, using FIMO predictions,^49^ we can identify >10,000 potential sites with the SFE motif across the genome (Table S7), suggesting the possible existence of many more SFEs.

**Figure 7.**
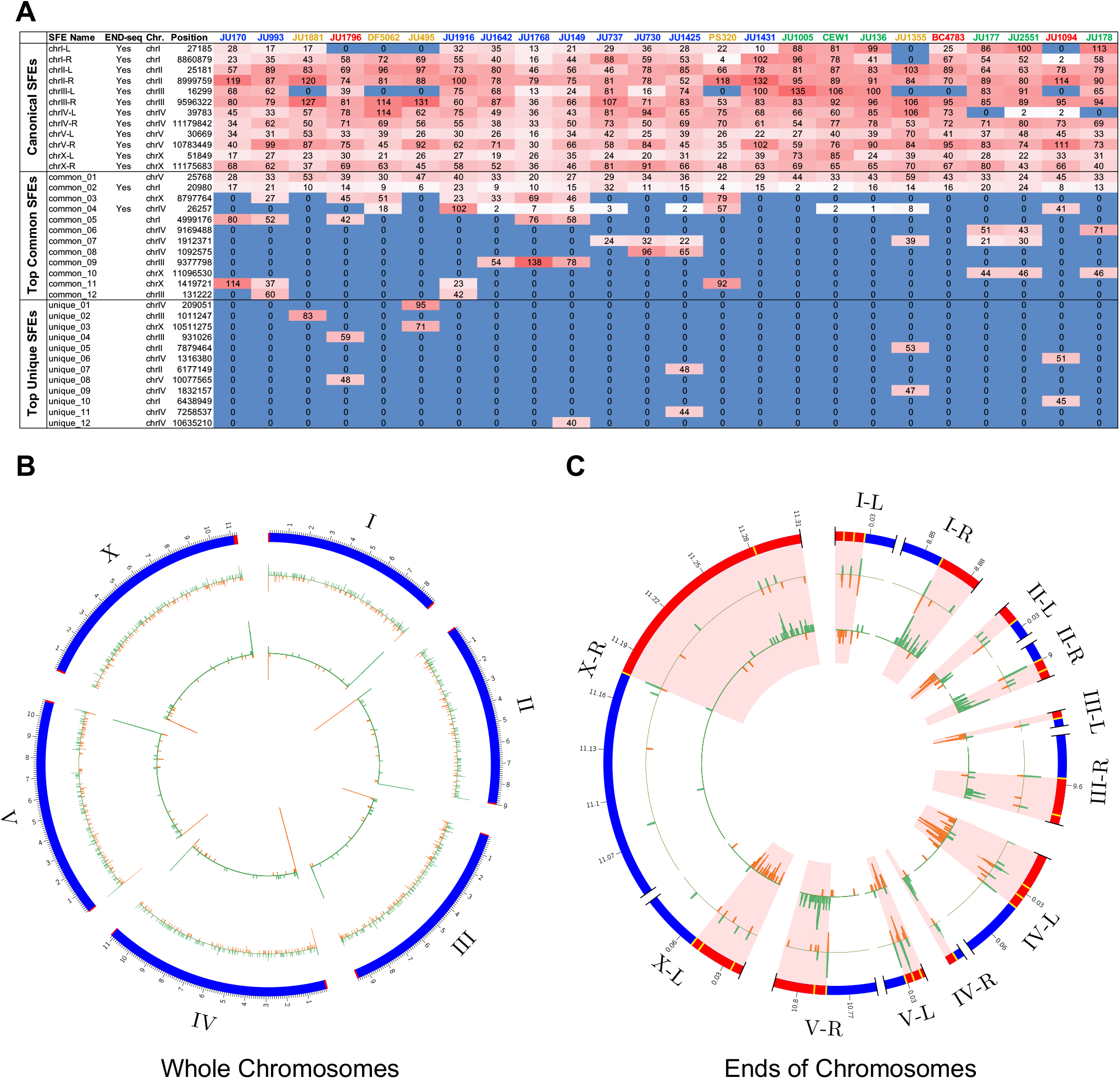
Widespread distribution of potential break sites at the ends of CEW1 chromosomes. **A**. Many new functional SFEs from the wild isolates can be found in CEW1 genome. The CEW1 chromosome coordinates and the normalized telomere-unique reads (out of 10 million reads) from the 24 *O. tipulae* strains are shown in a heatmap. The 12 canonical SFEs, top 12 common SFEs (seen in more than one strains) and top 12 unique SFEs (see in only one strain) are shown. Some canonical SFEs, including chrI-L, chrIII-L, and chrIII-R, do not have telomere reads in several wild strains. Additional SFE sites (399 in total) are in Table S6. **B** and **C**. Distribution of the SFEs in the CEW1 genome. Circos plot of CEW1 genome with FIMO predicted sites (score > 60; outside circle with random distribution across the genome, besides an enrichment at the chromosome ends; see Table S7 for the full list of sites) and the mapped 399 SFEs from the wild isolates (inside circle with highly biased distribution towards the ends). Green denotes sites on the forward strand and orange shows sites on the reverse strand. Retained DNA are blue and eliminated regions are red. Genomic coordinates are displayed in Mb. **B** shows the whole chromosome view, while **C** illustrates a zoomed in view of the eliminated ends (red shaded area; most of the retained DNA is not shown). The canonical and alternative SFEs are indicated in yellow in the outer circle. The majority of the mapped SFEs within the CEW1 eliminated regions (232/241, 96%) have the same strand as used in the wild isolates.

The distribution of these 399 SFEs is highly enriched at the ends of the CEW1 chromosomes (Fig. 7B-C). Most of them are in the eliminated regions, close to the canonical break sites, except for chrX-R, where most SFEs are within 30 kb at the end of the chromosome. In addition, the orientation of these SFEs is consistent with the nearby canonical SFEs, and most have the same conserved motif. The FIMO predicted sites using the conserved motif showed enrichment at the end of chromosomes, but additional predicted sites are seen throughout the genome, suggesting that sequence alone is not enough to functionally define the sites used for PDE (Fig. 7B-C). Collectively, our data illustrated that the *O. tipulae* CEW1 genome is organized to fulfill PDE. The data suggest that many potential SFEs are arranged adjacent to each other to ensure that DNA breaks occur at the chromosome ends for PDE.

## DISCUSSION

Programmed DNA elimination was first described by Boveri in the 1880s, yet the molecular details, mechanistic underpinnings, and functional importance of PDE are largely unknown, particular in multicellular organisms.^5-9^ Functional studies on metazoa PDE are lacking, largely due to the absence of a tractable model, where animals with mutations that alter the PDE process can be analyzed. In this study, we illustrate that *O. tipulae*, a free-living nematode with a 60-Mb genome, can be used as a functional and genetic model to study PDE.

Through analyses of staged embryos, we have determined that PDE occurs during early embryogenesis at the 2-16 cell stages. We also defined and analyzed the expression of the eliminated genes and features associated with the DNA breaks and telomere healing. A novel discovery for metazoa PDE is the identification of a sequence motif, the Sequence For Elimination (SFE), and its direct role in PDE. Our END-seq revealed that in the CEW1 strain, multiple SFEs, 12 canonical and 12 alternative, are used simultaneously at the onset of PDE. Additional data from the SFE mutants and wild isolates illustrates that many alternative SFE sites exist and can be used to ensure chromosome end removal and remodeling.^22^ The concerted and redundant mechanism for chromosome end remodeling leads us to suggest PDE likely serves an important function in *O. tipulae*.

### A comparison of PDE between the nematodes *O. tipulae* and *Ascaris*

The parasitic nematode *Ascaris* is one of the best-studied metazoan models for PDE.^5,6^ Our study allows for a direct comparison of PDE between *O. tipulae* and *Ascaris*. In both species, PDE occurs during early embryogenesis (2-16 cell stages). The ends of all chromosomes are broken, resulting in the loss of subtelomeric and telomeric sequences, and new telomeres are added to the broken DNA ends. In addition, PDE provides a means of DNA silencing in the somatic cells for both species by removing genes and repetitive sequences.^20-22^ Recently, PDE has also been discovered in other free-living nematodes from Clade V, including other *Oscheius* species and some *Caenorhabditis* and *Auanema* species [Gonzalez de la Rosa, Stevens, Pires-daSilva and Blaxter, personal communications], and *Mesorhabditis*.^50^ Other parasitic nematodes (Clade III *Parascaris* and *Toxocara*,^21^ *Baylisascaris*,^51^ and Clade IV *Strongyloides*^52^) also exhibit DNA break and sequence loss, illustrating the broad phylogenetic distribution of PDE in nematodes. However, other well-studied nematodes, including *C. elegans*, do not undergo PDE.^53^ Many nematodes have holocentric chromosomes with functional centromeres distributed along the length of the chromosomes,^54^ a feature that in principle allows DNA breakage to be more tolerable.^55,56^ Holocentric chromosomes also enable centromere reorganization that contributes to the selective segregation of retained DNA during *Ascaris* PDE.^30^ Do holocentric chromosomes render nematodes more adaptive to PDE? Another open question is whether basal nematodes had PDE or whether PDE has evolved several times independently in divergent nematodes.

There are notable differences in PDE between *O. tipulae* and *Ascaris. O. tipulae* exhibits no internal chromosome breaks that lead to changes of the number of chromosomes, while many *Ascaris* chromosomes have internal chromosome breaks that lead to an increase in its chromosomes from 24 in the germline to 36 in the soma.^22^ A second key difference is that *O. tipulae* break (SFE) sites have a distinct motif that is essential for PDE (Fig. 3), while *Ascaris* break regions appear to have no common sequence.^21^ The most conserved sequence within the *O. tipulae* SFE motif is the GGC/GCC, which likely provides a primer for telomere (TTA<u>GGC</u>) addition. In *Ascaris*, only one nucleotide of homology is needed to prime for *de novo* telomere addition.^21^ Thus, the difference in the conservation of sequence for the DNA breaks in the two species could be in part an adaption to a sequence requirement (or the lack of) for telomere addition in *O. tipulae*. Finally, the CBRs are associated with open chromatin during PDE in *Ascaris*. They remain open after PDE (at 32–256 cells), suggesting dynamic nucleosome or epigenetic factors are involved.^21^ In contrast, *O. tipulae* break regions are not associated with more accessible chromatin and do not show specific accessible changes; they appear to be relatively inaccessible (Fig. S9). This suggests that either the chromatin at the break sites is not required to be widely accessible for the break and repair or that the chromatin accessibility is transient in *O. tipulae*. Overall, the differences between *O. tipulae* and *Ascaris* provide a framework to study the variations in PDE mechanisms in both nematodes.

A final difference is that *O. tipulae* eliminates only a very small portion of its genome (0.6% = 349 kb, 112 genes), while *Ascaris* eliminates 18% of its genome (55 Mb, 918 genes).Furthermore, the expression profiles and functional annotation for the eliminated genes have notable differences. *Ascaris* eliminates many genes associated with spermatogenesis^20-22^ while the eliminated genes in *O. tipulae* are not enriched in males. Interestingly, several genes in *O. tipulae* appear to have a burst of expression only in the two primordial germ cells (PGCs) during embryogenesis (Fig. 2D). Considering the PGCs are regarded as transcriptionally quiescent in *C. elegans*,^57^ it is intriguing to see such a burst of expression for these eliminated genes in *O. tipulae* PGCs. Future studies are required to determine the function of these genes in the PGCs.

Our data indicate that PDE in *O. tipulae* likely occurs during early embryogenesis at the 2-16 cell stages. This is based on the observations from the genome sequencing through staged embryos (Fig. 1 and Fig. S2), imaging analyses of the presumptive elimination DNA foci (Fig. S3), and END-seq data on the timing of DNA break and telomere healing (Fig. 4E and Fig. S7). However, these methods have caveats. For genome sequencing and END-seq, although the staged embryos are synchronous, variations for the cell numbers in the embryos exist (see Fig. S1). In addition, the eliminated DNA residual following PDE mitosis may still be present in the cytoplasm for a few cell cycles. In comparison, the staining data can provide information on individual cells and embryos. However, due to the short cell cycle and the small amount of eliminated DNA, it may be challenging to observe DNA elimination in *O. tipulae* as has been seen in *Ascaris*, where the eliminated DNA remains at the metaphase plate during anaphase.^22^ It is also possible that the observed presumptive eliminated DNA foci in *O. tipulae* (Fig. S3) could come from the breakdown of the polar bodies as demonstrated in *C. elegans*.^58^ Thus, more precise and/or sensitive methods, such as better staging of embryos, single cell genomics, and live-cell imaging may be needed to further determine the exact stages of PDE in *O. tipulae*.

### A fail-safe mechanism for *O. tipulae* PDE

There are 12 canonical SFE sites at the junction of the retained and eliminated DNA on each chromosome that determine the sequences to be eliminated (Fig. 1 and Fig. 4). Surprisingly, our END-seq revealed 12 additional, alternative SFEs that reside in the eliminated regions (Fig. 5).

These 24 SFEs are used simultaneously at the onset of PDE (Fig. 5). Our mutants with disrupted canonical SFEs confirm that these alternative sites are used (Fig. 3D). Although not all eliminated regions contain an alternative site, our data suggest these alternative sites may serve as a fail-safe mechanism for *O. tipulae* PDE. In addition, genome sequencing and analyses on 23 wild isolates identified ∼400 possible alternative SFEs that tend to map to the ends of CEW1 chromosomes (Fig. 7). This suggests an evolutionary selection of specific sequences at the ends of the chromosomes in the *O. tipulae* genome to ensure the removal of the chromosome ends during PDE.

A key question in the field is the function and biological significance of PDE. Our *sfe-2l-Df61* mutant showed that failure to eliminate a single end of this chromosome led to no visible defects in the worms. However, this mutant only retains 10 kb of unique sequence. In comparison, the *sfe-xr-Df6* and *sfe-xr-In22* mutants failed to eliminate >100 kb of DNA (over 30% of total eliminated DNA in *O. tipulae*) but nevertheless removed the right end of the X chromosome. These mutants also showed no obvious detrimental phenotype, although further careful analyses might reveal subtle phenotypes. It is also possible that retention of multiple chromosome ends could uncover more deleterious phenotypes. We note that the mutants were cultured in a lab environment that may not mimic the challenges that exist in nature.Nevertheless, future work on mutants that fail to eliminate the full end of other single sites, multiple sites, or all ends of chromosomes promises to reveal novel insights into the function of PDE in *O. tipulae*.

Why does CEW1 only make breaks at these 24 sites out of the ∼400 potential SFEs? How are these sites selected? Once chosen, how are the breaks made and processed before telomere addition? We hypothesize that the conserved motif sequence (GGC/GCC) may play an essential role during the PDE process. Our SFE mutants showed that disruption of these sequences leads to a fail-to-eliminate phenotype (Fig. 3). The SFE is a degenerate palindromic sequence that could be recognized by a DNA-binding protein (Fig. 3 and 5). Thus, one possibility is that a DNA-binding protein recognizes these sites during PDE. Once bound, the protein can generate the DNA breaks directly or by recruiting other proteins to the sites. In these scenarios, additional factors would be necessary to distinguish these key 24 sites from other potential non-functional sites. These factors could be related to the binding affinity of the DNA/protein interactions.

Additional factors might include the 3D genome organization, and/or other epigenetic features. It is also possible that the recognition of the break sites is mediated by cis- and trans-regulatory RNAs (small RNAs or lncRNAs) or other epigenetic factors without the involvement of a specific DNA-binding protein.^59^ Alternatively, DNA replication stress, RNA transcription, and R-loops^60-62^ induced by the SFEs may contribute to the generation of the DNA breaks. Surprisingly, the chrX-L mutant (*sfe-xl-Df106*) used a novel site nearby (473 bp) instead of the two existing alternative SFE sites in response to a disrupted canonical SFE site (Fig. 3D and 5B). This indicates that this novel site has the potential to be a break site, but in the wild type, the use of the adjacent canonical break prevents it from serving as an SFE.

In addition to the usage of a novel break site, several other interesting features are associated with the *sfe-xl-Df106* mutant. First, the sequence coverage for the eliminated region at the left end of the X chromosome is uneven and in general higher than other eliminated regions (29.8% vs. 23.6%). Also, the novel break site has a low FIMO score (47, ranked 5257) and it is not an alternative site from the wild isolates or from the END-seq data (Fig. 7). One possibility is that multiple alternative break sites could be used in independent PDE events (in each pre-somatic cell) in this mutant, causing the unusual read coverage (Fig. 3D). Additionally, in this mutant, two genomic regions at the ends of chromosomes (chrI-R and chrX-R) within the eliminated DNA have an unexpected high somatic read coverage (Fig. 3D, blue arrow). This high copy number suggests these sequences may have been duplicated and inserted to the retained regions of the genome. This sequence duplication and rearrangement is reminiscent of genome instability often seen in pathological conditions (such as aging and cancers).^63,64^ Interestingly, these two regions encode genes that are believed to undergo a burst of transcriptional expression in the PGCs of the wild type CEW1 (Fig. 2D). How could a deletion of this SFE site cause genome instability? These features of the *sfe-xl-Df106* mutant merit future in-depth investigation.

## CONCLUSIONS

PDE contradicts the genome constancy rule, yet it is seen in many phylogenetically divergent groups, including multiple metazoa, suggesting its broad molecular and biological significance. The molecular mechanisms of PDE in multicellular organisms remain largely unknown, partly due to the challenges of working with organisms that are not traditional models and have limited tools. Here we present a new model to study PDE in the free-living nematode *O. tipulae*. Our genomic and functional data reveals the presence of a conserved motif (SFE) that directly facilitates PDE, a feature that has yet to be described for other metazoa PDE models. Our genetic and molecular analyses show that alternative SFEs can be utilized when necessary, perhaps serving as a fail-safe mechanism for PDE. The amenability to genetic and functional manipulations, short life cycle, modest amount of DNA eliminated, and the ability to capture discrete embryonic stages, including those stages that undergo PDE, makes *O. tipulae* an excellent model to carry out in-depth molecular studies that will uncover the functions, mechanisms, and consequences of PDE in a multicellular organism.

## MATERIALS AND METHODS

### Worm strains, culture and maintenance

The CEW1 strain (originally isolated by Carlos E. Winter in Brazil) and the HIM-1 mutant (PS2626) were obtained from CGC (https://cgc.umn.edu/). The wild isolates of *O. tipulae* were from Dr. Marie-Anne Félix. The maintenance of *O. tipulae* is essentially the same as *C. elegans*. Briefly, *O. tipulae* strain CEW1, HIM-1 and other wild isolates were cultured using standard Nematode Growth Medium (NGM) plates seeded with *E. coli* OP50 bacteria.^65^ To prepare a large number of synchronous worms, we used the *C. elegans* liquid culture protocol^66^ with adaptations to promote dauer formation. Dauer worms were isolated using a 1% SDS solution followed by sucrose float. They were immediately plated on rich agar media containing an *E. coli* NA22 bacterial lawn. The dauer develop and mature into egg-laying worms at room temperature within ∼36-40 hrs.

### Egg isolation, staged embryos and larvae, and other samples

Egg-laying worms were washed off plates and passed through a 70-µm mesh. This step allows existing eggs to fall through the mesh, leaving the mature worms on the top. These worms were then bleached with sodium hypochlorite solution (Sigma Cat# 425044-250ML, 0.5 M NaOH, 3% bleach) to collect embryos (largely 2-4 cell stage, Fig. S1). Early embryos were further developed in Virgin S Basal (VSB) medium (100 mM NaCl, 5.7 mM K^2^HPO4, 44.1 mM KH^2^PO^4^) at 25°C to reach to the desired embryonic stages (Fig. S1). Briefly, embryos were staged by examining samples at half-hour intervals using a Leica DMIL-LED microscope with a 10X 0.25 NA lens and a Leica DMC6200 camera. Images were scored by counting the number of cells visible in each embryo to determine the stage.

To prepare larval stages, the flow-through of the 70-µm mesh was further passed through a 25-µm mesh to separate mixed embryos from various stages of worms. These mixed embryos were developed in the VSB medium until all embryos reach to L1. The arrested L1s were plated on NGM with a robust OP50 lawn to develop at 20°C to synchronous stages of larvae, young adults, and mature adults. Alternatively, post-dauer larvae and worm samples were collected starting from the dauer harvested from the liquid culture. For adult males, we handpicked 200 males from the HIM-1 mutant for RNA-seq library replicate.

### Hoechst and immuno-staining

The staining procedure was adapted from a previous study on *Ascaris* embryos.^67^ Briefly, embryos obtained from bleached adults were spun down at 1,000 x g for 2 minutes in a 1.5 mL Eppendorf tube and resuspended in 500 µL of fixative solution containing 50% methanol (Fisher Chemical, A412SK-4) and 2% para-formaldehyde (Ted Pella, Inc., Cat # 18505) in 1X PBS buffer. The embryos were freeze-cracked by submerging the tube in liquid nitrogen until the solution was completely frozen, followed by thawing in a beaker of lukewarm water. This process was repeated at least five times to ensure thorough eggshell cracking. The tube was opened between each round to release trapped nitrogen gas. After spinning and removing the fixative, the embryos were rehydrated by incubation in 25% methanol in 1X PBS for one minute, followed by incubation in 1X PBS for one minute. Embryos were resuspended in 1 mL of blocking solution (0.5% BSA in 1X PBS) and incubated for 30 minutes at room temperature with rotation. Embryos were then spun down and resuspended in blocking solution and stained with primary antibody (mostly diluted 1:1000) overnight at 4°C with rotation. Secondary antibody incubation (diluted 1:1000) was carried out at room temperature for two hours in the dark with rotation, followed by washing in blocking solution. Embryos were then incubated with 500 µL of Hoechst 33342 (1 µg/mL) (Invitrogen, Fisher Cat# H3570) dye at room temperature for ten minutes in the dark with rotation. Embryos were washed in blocking solution and mixed with Vectashield plus antifade mounting media (Vector Laboratories, H-1900-10) and pipetted onto a glass slide. For each slide, a coverslip was placed on the sample and the slide was wrapped in a paper towel, turned upside-down, and gently flattened with a large book for 3 minutes.Coverslips were sealed using clear nail polish (Ted Pella, Inc., Fisher Cat# NC1849418). Imaging was performed on a Nikon Eclipse Ti inverted spinning disk confocal microscope using a 100X 1.49 NA oil immersion objective lens. Images were collected on a Photometrics Evolve 512 EMCCD camera and acquired using MetaMorph software (version 7.7.10.0, Molecular Devices, LLC). Images were acquired as Z-stacks using a step size of 0.3 µm. Image processing was performed using FIJI.^68^ All contrast adjustments in figures are linear.

### RNA isolation and sequencing

Total RNA was prepared using TRIzol (Invitrogen Cat# 15596026) protocol. The quality of the RNA was evaluated with the TapeStation 4200 (Agilent) and quantified with the Qubit(tm) 4 Fluorometer (Thermo Fisher). The ribosomal RNAs were removed using the RiboCop rRNA Depletion Kit (LEXOGEN Cat# No 144). The RNA-seq libraries were constructed using the CORALL Total RNA-Seq Library Prep Kit (LEXOGEN Cat# No 146) and sequenced using an Illumina NovaSeq 6000 at the University of Colorado Anschutz Medical Campus Genomics Core.

### Genomic DNA extraction and sequencing

Cultures of *O. tipulae* were grown on 150 mm plates of rich agar seeded with *E. coli* NA22. When the bacterial food source was near exhaustion, worms were washed off plates with M9 buffer and centrifuged at 170 x g for 1 min. The pelleted worms were then purified from microbial contaminants using a 35% sucrose floatation.^69^ Genomic DNA was extracted from harvested worms using Genomic-tips (Qiagen Cat# 10223). The resulting high molecular weight genomic DNA was used to prepare sequencing libraries using the Illumina DNA Prep kit (Cat# 20018704) following the manufacturer’s instructions. The libraries were sequenced using an Illumina NovaSeq 6000. The 23 wild isolates were sequenced using the same procedure.

### CRISPR-Cas9 modification of the *O. tipulae* genome

We adapted CRISPR-Cas9 procedures from *C. elegans*^70,71^ and other nematodes^72^ and developed a protocol for *O. tipulae*. All CRISPR-Cas9 reagents (crRNAs, tracrRNAs, Cas9, Cat# 1081058) were obtained from IDT DNA (Coralville, IA). Locus-specific crRNAs were selected using a combination of rating algorithms from IDT DNA and CRISPRscan.^73^ Candidate crRNAs that scored at least 40 (range 0-100) in both algorithms were considered for further experimentation. A list of all crRNAs and repair templates used is provided in Table S1. The injection mixes were synthesized as follows: A CRISPR-Cas9 RNP to induce a dominant roller mutation in *O. tipulae rol-6* (see below) for co-CRISPR selection was made by mixing 0.5 µL 10 mg/mL Cas9 protein into 4.5 µL IDT duplex buffer. 2.5 µL of 100 µM tracrRNA was added, followed by 2.5 µL of 100 µM *rol-6* crRNA. A locus-specific CRISPR-Cas9 RNP was produced by mixing 0.5 µL 10 mg/mL Cas9 protein into 3.5 µL IDT duplex buffer. We then added 4.0 µL of 100 µM tracrRNA, followed by 1.0 µL of 200 µM of each crRNA that flanked the elimination motif. Both CRISPR-Cas9 RNP complexes were incubated at 37°C for 15 minutes, then 1.0 µL of 100 µM repair template was added to each RNP mix to mediate homology-directed repair of a roller phenotype or mutation of an SFE motif. The two RNP complexes were mixed and centrifuged at 10,000 x g for two minutes. Lipofectamine (Invitrogen Cat# 13778030) was added to 3% v/v to the RNP-containing supernatant, and the mix was incubated at room temperature for 20 minutes prior to loading needles for injection.

Injections were done using a Nikon Eclipse Ti-U inverted scope with DIC optics and a Narishige IM 300 injector with an input pressure of 70 psi and an injection pressure of 20 psi. Injection needles were from World Precision Instruments Inc. (cat. # 1B100F-4) and were pulled on a Sutter P-87 puller using a two-step cycle of heat 330, pull 0, velocity 20, time 200, followed by heat 330, pull 20, velocity 65, time 150. The pressure was set to 400.

Well-fed young adult (4 days old) *O. tipulae* were picked to pads of dried 2% agarose and covered with 700 weight halocarbon oil (Sigma Cat# 9002839) for injection. Injected worms were released from agarose pads by gentle resuspension in M9 buffer, picked to OP50-seeded NGM plates, and incubated at 25°C. Plates were screened for F1 worms with a roller phenotype at days 3-5 post-injection. In our hands, as many as 15% of injected P0 worms produced one or more F1 progeny with a roller phenotype. F1 rollers were allowed to produce F2 progeny, and either the parental F1 roller or a pool of 10 F2 progeny were screened by PCR ^70^ to detect the CRISPR-Cas9-induced edit of interest. Potential mutants were plated as single F2s, and their progeny were screened for homozygosity for the mutation of interest via PCR.

### Identification of a *rol-6* orthologue in *O. tipulae*

Certain alleles of the *C. elegans rol-6* locus, which encodes a cuticular collagen, give rise to a dominant “roller” phenotype^74^ that is readily scored and can be used for co-CRISPR selection. Orthologous *rol-6* genes have also been identified in *Pristionchus pacificus* and *Auanema* species ^72,75^. Using the *Auanema rhodensis* ROL-6 protein sequence, we scanned the *O. tipulae* genome and identified a candidate *O. tipulae rol-6* gene (Fig. S10), encoding a protein that has 82% identity through an 88-amino acid segment of *A. rhodensis* ROL-6 that encompasses the critical R→C substitution associated with a roller phenotype (Fig. S10). The introduction of the R→C substitution via CRISPR-Cas9 and a repair template harboring the desired base substitutions (Fig. S10) resulted in *O. tipulae* worms with a roller phenotype essentially indistinguishable from that of *C. elegans* with a *su1006* allele of the *rol-6* gene. This mutant *rol-6* repair template was used in all subsequent experiments as a co-CRISPR selection marker.

### END-seq library preparation

Staged embryos were used to create END-seq libraries using an adapted protocol.^47,48^ Briefly, embryos were embedded in agarose plugs to protect the DNA from exogenous breaks. Some plugs were digested with the restriction enzyme AsiSI (NEB, catalog # R0630) or FseI (NEB, catalog # R0588) to generate DSBs as an internal control. DSBs were blunted with exonuclease VII (NEB, catalog # R0630) and exonuclease T (NEB, catalog # M0625). Blunt ends were A-tailed and capped with END-seq adaptor 1, a biotinylated hairpin adaptor. DNA was liberated from the plugs and sheared to ∼200-300 bp with a Covaris M220 focused-ultrasonicator (130 µL tube, peak power 50, duty 16, cycles/burst 200 for 420 seconds). DNA fragments containing END-seq adaptor 1 were isolated with Dynabeads^™^ MyOne^™^ Streptavidin C1 (Invitrogen, catalog # 65001). The other ends broken by sonication were repaired and A-tailed with END-seq adaptor 2. Hairpins were digested with USER (NEB, catalog # M5505), and the DNA was amplified with Illumina TruSeq primers and barcodes. The libraries were sequenced with an Illumina NovaSeq 6000.

### ATAC-seq library preparation

Staged embryos were used to carry out ATAC-seq. Embryos were resuspended in 5 mL of cold ATAC-seq lysis buffer (10 mM Tris-HCl pH 7.4, 10 mM NaCl, 3 mM MgCl2, 0.1% Triton-X 100, 0.1% NP40 alternative) and transferred to a dounce homogenizer kept cold on ice. Embryos were dounced ten times before being transferred to a 15 mL Eppendorf tube and spun down at 500 x g for 5 minutes at 4°C. Samples were tagmented and subsequently purified using an ATAC-seq kit (Active Motif, 53150) according to the cell sample protocol. Libraries were amplified through ten rounds of PCR and sequenced using an Illumina NovaSeq 6000.

### Bioinformatic data analysis

#### Genome mapping

Genomic reads from staged embryos, SFE mutants, and wild isolates were mapped to the *O. tipulae* telomere-to-telomere genome^34^ using bowtie2^76^ and SAMtools^77^ to generate BAM files. The read coverage was obtained using BEDTools^78^ to produce a BEDGRAPH file that was loaded into UCSC genome browser track data hubs.^79^

#### RNA-seq data analysis

Ribosomal RNA reads were filtered out using Bowtie2^76^. The non-rRNA reads from each sample were mapped to the *O. tipulae* genome with STAR.^80^ The transcripts for each sample were assembled separately using the StringTie^81^ and merged into a non-redundant set of transcripts. The predicted coding regions (https://github.com/TransDecoder) from the transcripts were used to further evaluated and to remove artifacts, cryptic transcripts, and transcriptional noises based on their coding capacity (< 50%), the lack of splicing, and/or low expression. For expression analysis, we used the HTSeq^82^ to get the read count for all transcripts. The DESeq2^83^ package was used to determine differentially expressed genes between the developmental stages with a fold-change cutoff of 2 and a p-value ≤0.05. Several R packages were also used to generate the heatmap,^84^ the volcano plot^85^ and the PCA plot (https://rdrr.io/cran/factoextra/).

#### Gene annotation and functional analysis

The transcripts were used to blast^86^ against protein databases (NCBI nr, UniProt,^87^ Swiss-Prot, WormBase^88^ *C. elegans* and *Ascaris*) to assign annotation. The translated protein sequences from TransDecoder were used to search against the Pfam database^89^ (hmmsearch E-value cutoff <=10^−10^). Genes that matched to *C. elegans* (WBgene ID) were used for Gene Ontology and tissue enrichment analysis^44,45^.

#### Motif analysis

*De novo* motif identification was initially done by analyzing 1 kb regions from the 12 chromosomal breakage sites in CEW1. These sequences were analyzed with MEME,^90^ a tool to discover ungapped motifs. *De novo* identification of the motif sequence was reinforced by including 23 wild stains in the analysis. The MEME motif matrix was used as input for FIMO^49^ to search for alternative sites within the CEW1 strain and among 23 wild isolates using default parameters (*p*-value < 1.0E-4). Telomeric sequences were filtered from the FIMO-generated motifs list, and a cut-off score of 50 was initially used to call the alternative motif sites.

#### END-seq data analysis

Reads (R1 file only) were mapped to the *O. tipulae* reference genome^34^ with Bowtie2^76^ and processed with SAMtools^77^. The 5’ position of each read was mapped, and its strand was determined with BEDTools^78^ (genomecov). For comparison among libraries, the reads were normalized to one million genome-mapped reads. Telomere-unique reads were defined as those containing two or more sequential telomeric repeats (TTAGGCTTAGGC).

Germline genomic regions that contain telomere repeats were filtered out. All telomere-unique reads were converted to the G-rich strand (TTAGGC). Cutadapt^91^ (-j 0 -m 20 -b TTAGGCTTAGGC) was then used to trim the telomeric sequences. These trimmed reads were mapped to the genome to identify the telomere addition sites.

#### ATAC-seq data analysis

ATAC-seq reads were mapped to the *Oscheius tipulae* genome using bowtie2^76^ and processed using SAMtools^77^ and BEDTools.^78^ The reads were normalized based on the genome-mapped reads. ATAC-seq data was uploaded to the UCSC genome browser track data hubs.^79^

#### Wild isolates sequence analysis

Raw reads were preprocessed with Trimmomatic^92^ to remove potential adapters and trim low-quality nucleotides. *De novo* genome assemblies of 23 wild strains were done using SPAdes^93^ with “--careful” option recommended for small genomes that reduces the number of mismatches and short indels. The mitochondrial genomes were assembled using a reference-assisted *de novo* approach as described.^94^ The phylogenetic tree for the mitochondrial genomes was constructed using MEGA11.^95^ The genomic reads were mapped to the CEW1 genome as described in the genome mapping section. Telomere-unique reads, described in the END-seq data analysis, were mapped to the CEW1 genome to identify potential sites of alternative SFEs.

## Abbreviations

PDE: programmed DNA elimination
SFE: sequence for elimination
CBR: chromosomal breakage region
DSB: DNA double-strand break
PGC: primordial germ cell
GO: Gene Ontology

## Data availability

All sequencing data will be deposited at the NCBI SRA and GEO databases - pending. The data for genome sequencing, gene models, RNA-seq, END-seq, and ATAC-seq are also available in a UCSC Genome Browser track data hubs^79^ that can be access with this link: http://genome.ucsc.edu/s/jianbinwang/CEW1-genome-browser.

## ACKNOWLEDGEMENT

We thank CGC for *O. tipulae* strains, Aurélien Richaud and Marie-Anne Félix for the wild isolates and sharing unpublished sequencing data, Pablo Manuel Gonzalez de la Rosa and Mark Blaxter for personal communications and sharing genomic data, Sally Adams and Andre Pires da Silva for liposome based CRISPR methods, and Chris Turpin, Jenny Heppert, Guy Caldwell, Laura Berkowitz, and Tom Evans for worm methods, protocols, and suggestions. We also thank graduate students Jansirani Srinivasan and Yingjie Xu and undergraduate students Eduardo Villalobos, Ryan Qiu, Mollie Sterling, Gracie Chiampas, Matthew Carr, and Jordan Parker for helping with worm maintenance, sample collection, and/or genetic analysis, the University of Colorado Anschutz Medical Campus Genomics Core for sequencing services, and Mariano Labrador, Albrecht Von Arnim and Dick Davis for comments and critical reading of the manuscript. This work was supported by NIH grant AI155588 and the University of Tennessee Knoxville Startup Funds.

## AUTHOR CONTRIBUTIONS

T.C.D. and J.W. designed the project; J.R. and J.W. developed protocols and prepared staged embryos; J.R.S. did staining and imaging; J.R. carried out RNA-seq; T.C.D. and V.T. performed CRISPR and genetic analyses; T.C.D. and E.S. sequenced genomes from staged embryos, mutants, and wild-isolates; B.E. did END-seq; J.R.S. did ATAC-seq; B.E., S.B.Z., M.V.Z., and J.W. carried out bioinformatic analyses; and T.C.D., B.E., and J.W. wrote the manuscript.

## List of Supplemental Figures

Figure S1. Staged embryos of *O. tipulae*.

Figure S2. Genome coverage and the timing of *O. tipulae* PDE during early embryogenesis. Figure S3. Hoechst and immuno-staining of *O. tipulae* embryos.

Figure S4. Improved *O. tipulae* gene models with RNA-seq data.

Figure S5. Differentially expressed genes in *O. tipulae* early embryos and males. Figure S6. CRISPR-Cas9 induced *sfe* mutations in *O. tipulae*.

Figure S7. Sites and frequencies for telomere addition in *O. tipulae*.

Figure S8. DNA break and telomere addition through *O. tipulae* early embryogenesis. Figure S9. ATAC-seq through *O. tipulae* early embryogenesis.

Figure S10. *O. tipulae* rol-6 gene, CRISPR, and its mutant phenotype.

## List of Supplemental Tables

Table S1. CRISPR RNAs, DNA repair templates, and PCR primers used in this study. Table S2. Stages and libraries for *O. tipulae* RNA-seq.

Table S3. Gene annotation and RNA expression for *O. tipulae*.

Table S4. Differentially expressed genes and GO enrichment between various *O. tipulae* stages. Table S5. Description of 23 wild isolations of *O. tipulae*.

Table S6. SFE sites identified from the wild isolates of *O. tipulae*.

Table S7. Potential SFE sites in *O. tipulae* CEW1 genome from FIMO prediction.

