## Supplemental Figures for "The Nematode *Oscheius tipulae* as A Genetic Model for Programmed DNA Elimination"

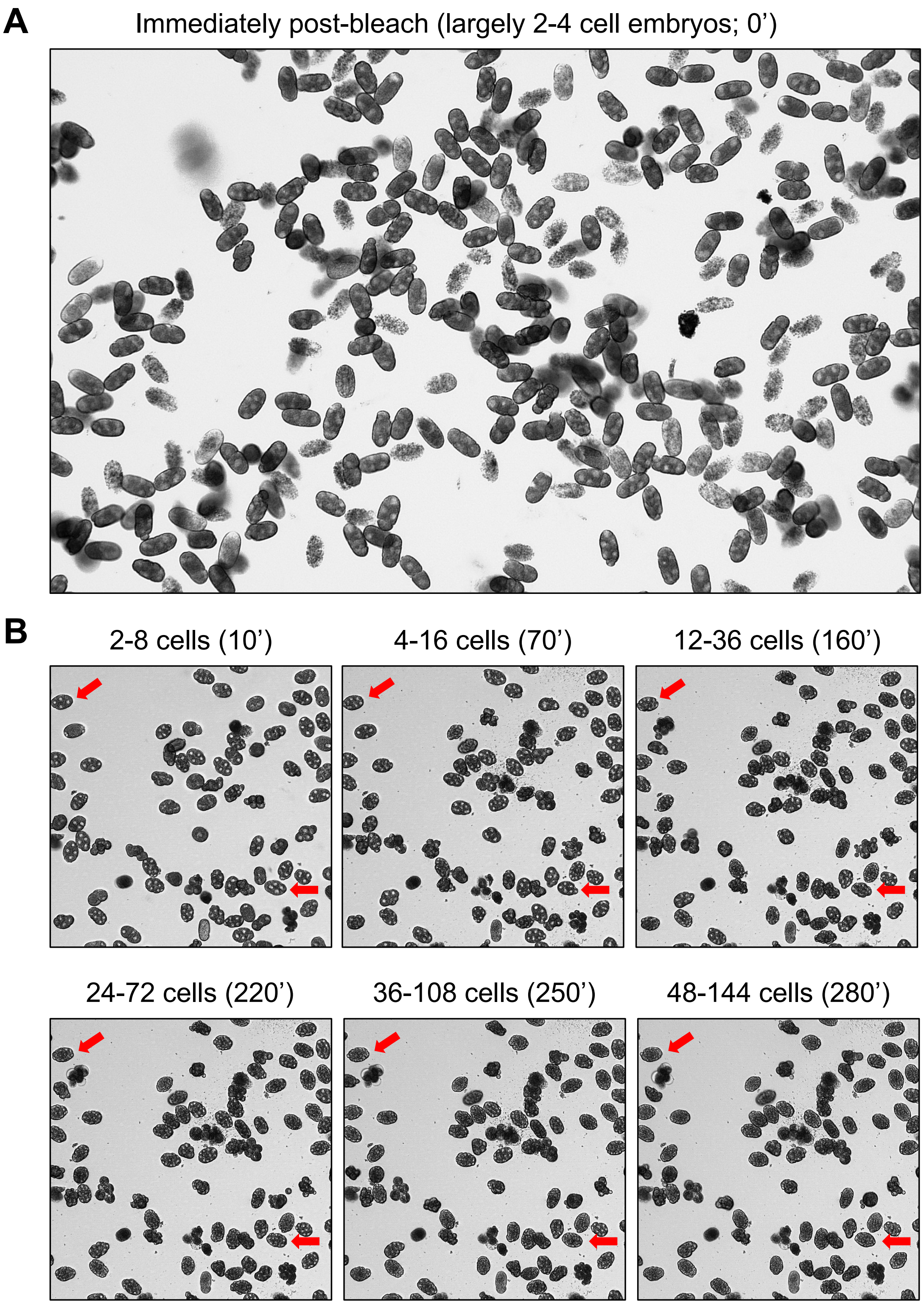


**Figure S1. Staged embryos of *O. tipulae*. A**. A representative image showing *O. tipulae* eggs immediately after harvest from adult worms. These eggs are designated as 0' embryos. **B.** 0' embryos were incubated at room temperature for the indicated time (minutes) to reach discrete embryonic stages. The same field of view is shown, with the red arrows tracing the development of two exemplary embryos.

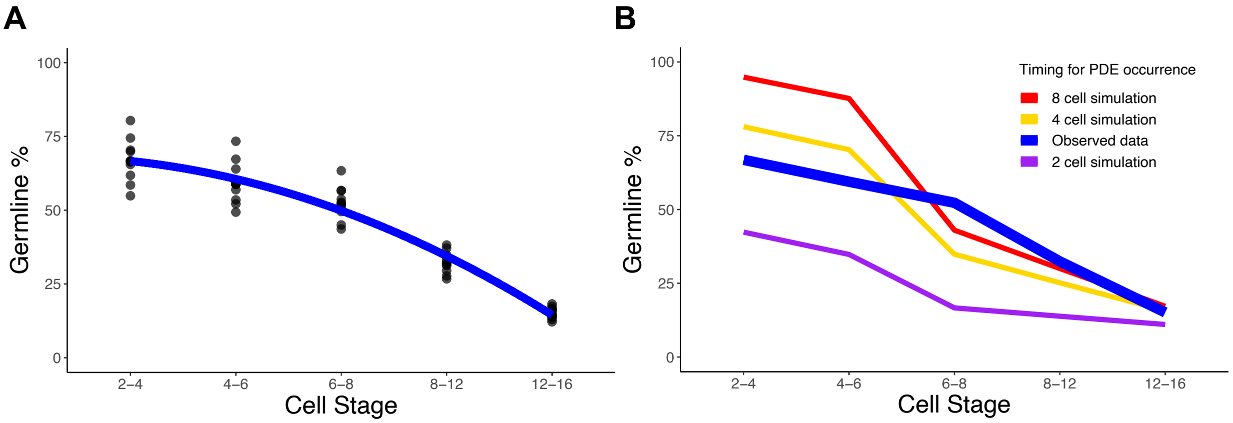


**Figure S2. Genome coverage and the timing of *O. tipulae* PDE during early embryogenesis. A.** The amount of germline DNA at the ends of chromosomes drops progressively as embryos develop. Shown are the ratios of read coverage for the 12 chromosome ends (each dot represents one end) over the average genome coverage for the retained DNA. The blue best-fit line (a second-degree polynomial) shows the overall trend of DNA loss through developmental stages. **B.** Comparison of genome coverage for the eliminated DNA between simulated and observed data. The simulation is based on counting the number of cells in representative staged embryos assuming that PDE begins at the 2-, 4-, or 8-cell stages. The trends for these simulated results do not agree with the observed data. The main discrepancy between these trends is at the 6-8 cell stage, where the simulation suggests a rapid drop in coverage while the sequencing data shows a gradual decrease. This discrepancy is likely due to the time required to fully degrade the eliminated DNA. The variations of the staged embryos used for sequencing may also contribute to the observed discrepancy. Considering these caveats and the low coverage (65%) observed at the 2-4 cell stage, we predict that *O. tipulae* PDE likely starts at the 2-cell stage.

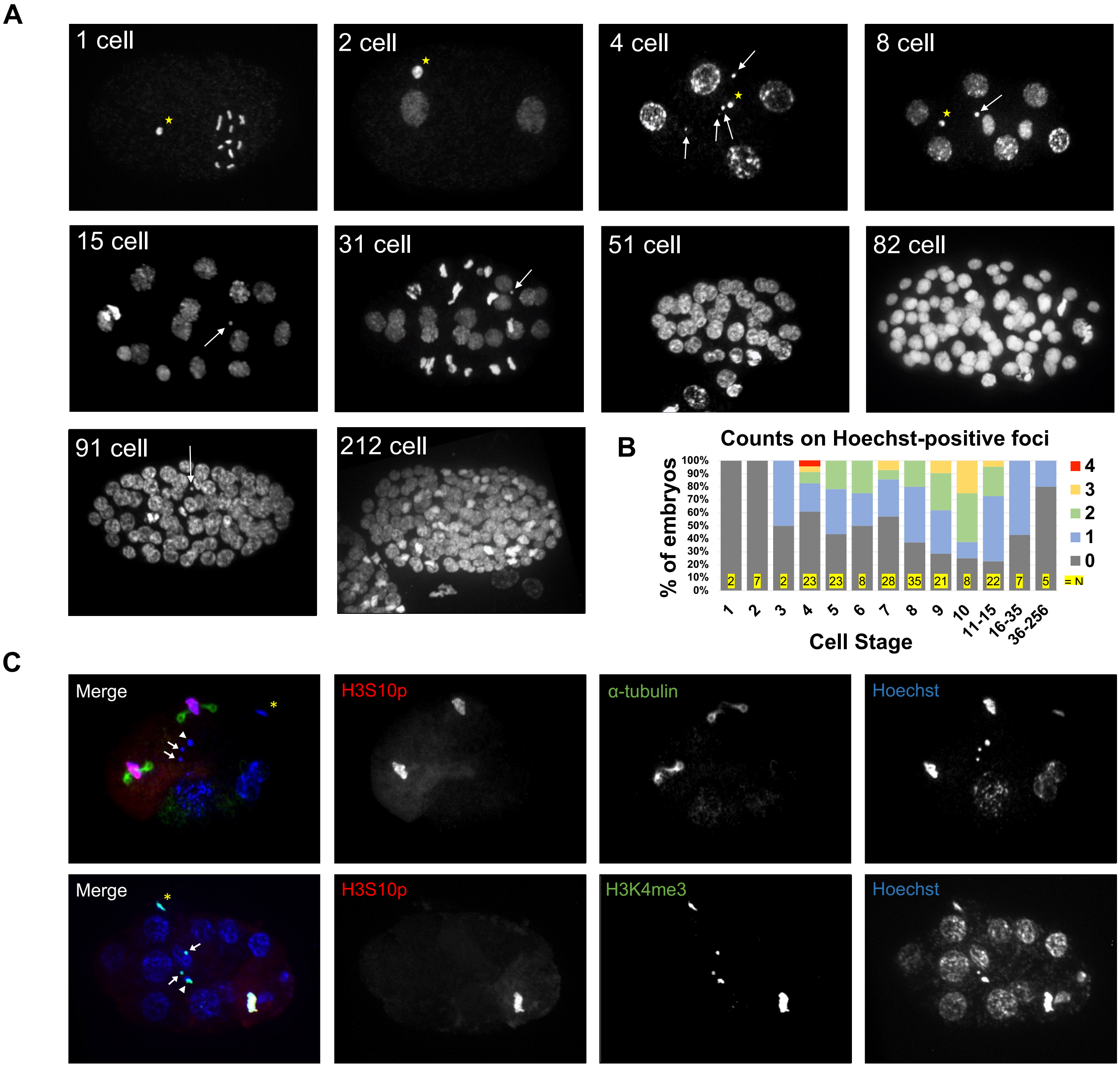


**Figure S3. Hoechst and immuno-staining of *O. tipulae* embryos. A.** Hoechst staining of *O. tipulae* embryos. The presumptive polar body is labeled with a yellow star, and the presumptive eliminated DNA is indicated with an arrow. **B.** Quantification of the presumptive eliminated DNA in *O. tipulae* embryos based on the number of DNA foci. A total of 191 embryos from different stages were examined. Note the Hoechst-positive foci are mostly seen during the 3-35 cell stages. **C.** Immuno-staining of early embryos. Hoechst-positive foci are shown with arrows in the merged image while polar bodies from the second meiotic division are shown with arrowheads and polar bodies from the first meiotic division are labelled with yellow asterisks. Top: A five-cell embryo where the ABa and ABp blastomeres are in metaphase, with the condensed chromosomes labeled by H3S10p and spindles stained by α-tubulin. The two Hoechst-positive foci (arrows) are not stained with H3S10p or α-tubulin. Bottom: A 14-cell embryo (not all cells are shown) with one cell in metaphase (bottom right) stained by H3S10p and H3K4me3. Two Hoechst-positive foci (arrows) are present and stained by H3K4me3 but not H3S10p. All images are displayed as Z-max projections.

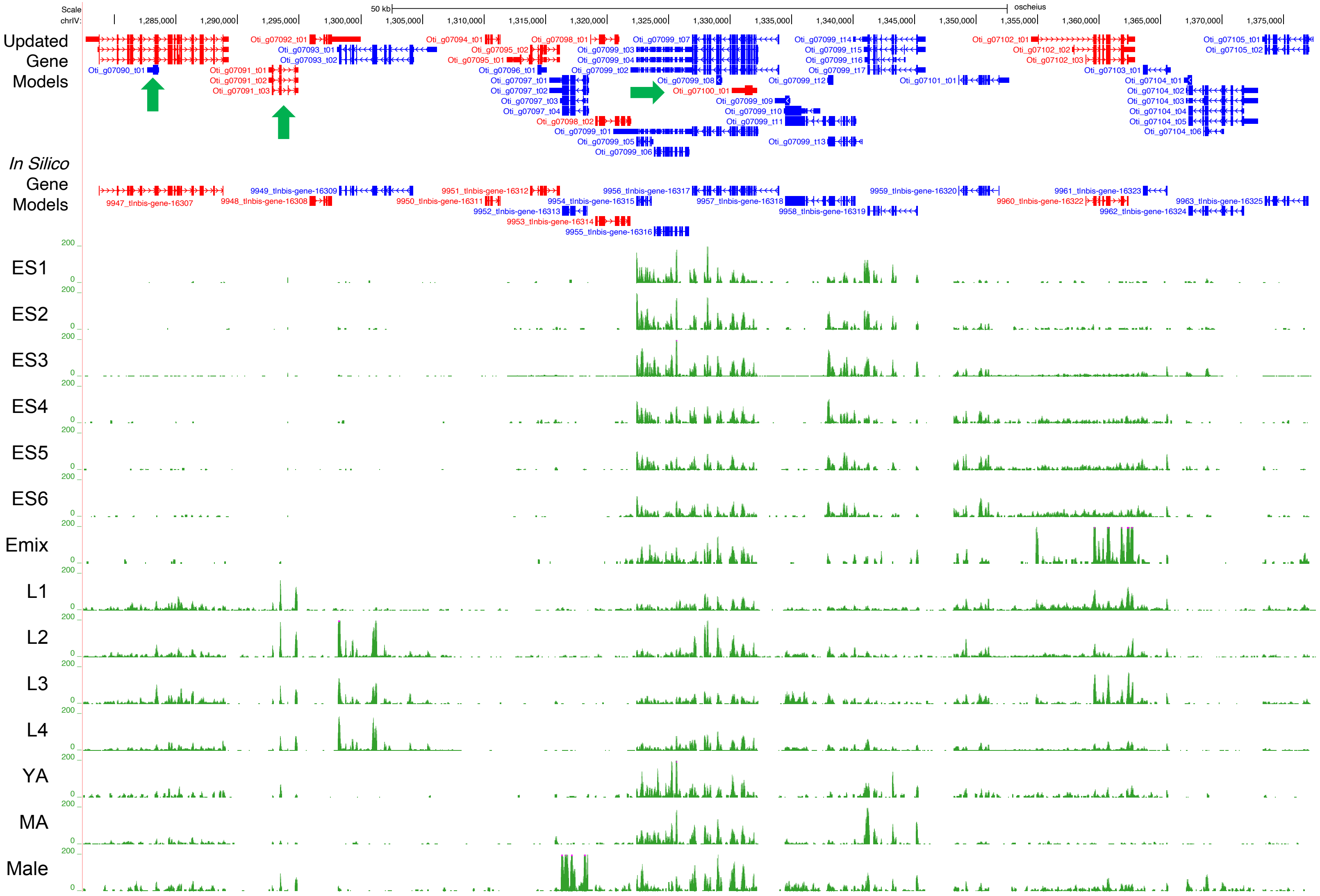


**Figure S4. Improved *O. tipulae* gene models with RNA-seq data.** Shown is a browser view of a 100 kb region from chromosome IV. For the gene models, RNA transcribed on the plus strand is red and on the minus strand is blue. Note the extensions on the 5' and 3' UTRs, the alternative isoforms, and the newly identified genes (green arrows). Normalized RNA-seq data for various stages are shown below in green tracks. The sample descriptions are: ES1 = 2-4 cells, ES2 = 4-8 cells, ES3 = 8-16 cells, ES4 = 16-32 cells, ES5 = 32-64 cells, ES6 = 64-128 cells, Emix = mixed embryos, YA = young adult, and MA = mature adult. The genome browser is available at http://genome.ucsc.edu/s/jianbinwang/CEW1-gene-models-RNA-seq.

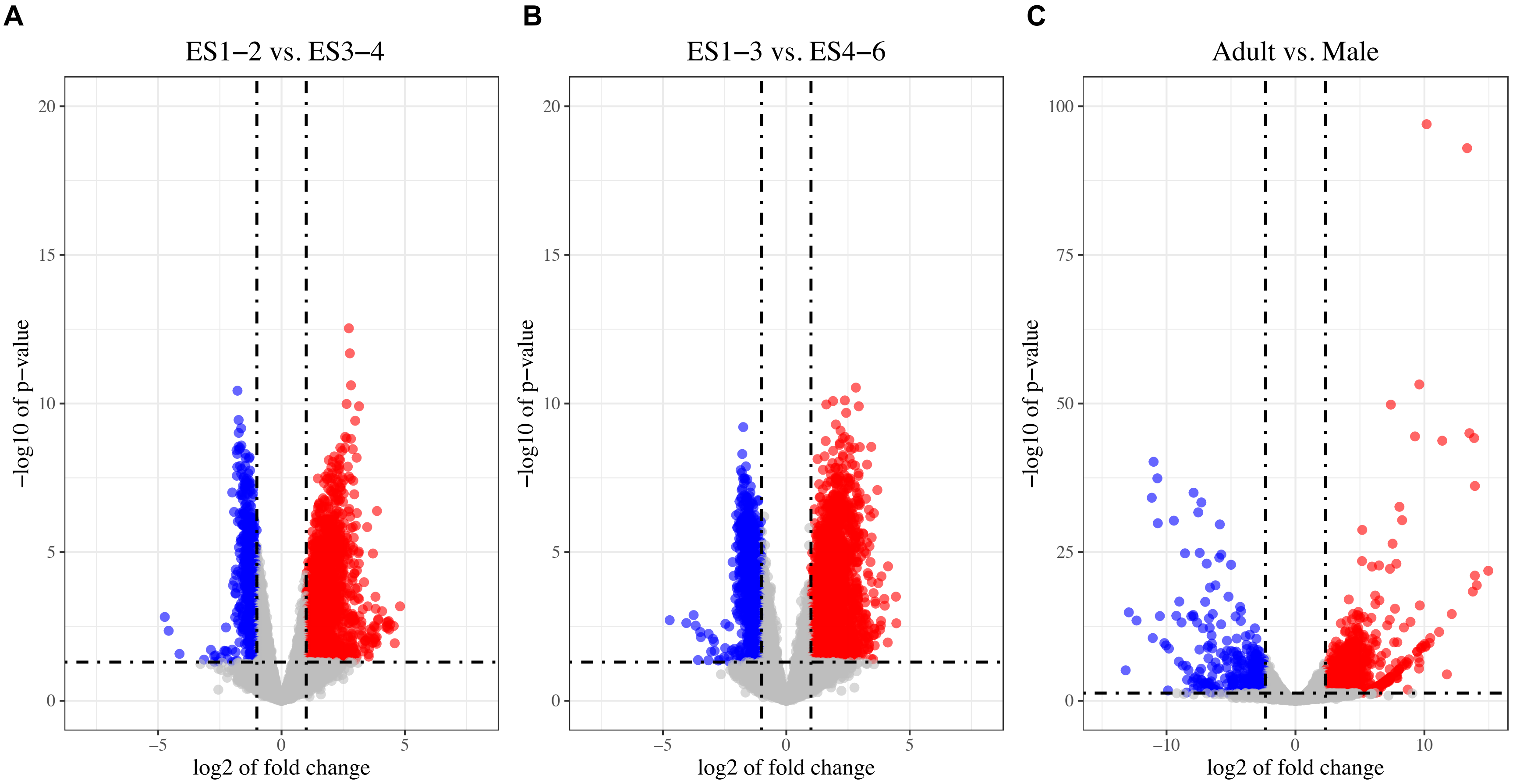


**Figure S5. Differentially expressed genes in *O. tipulae* early embryos and males.** Volcano plots showing differentially expressed genes in early embryos (**A** and **B**) and enriched genes in males (**C**). Sample descriptions: ES1-2 = 2-8 cells, ES3-4 = 8-32 cells, ES1-3 = 2-16 cells, and ES4-6 = 16-128 cells. Differentially expressed genes were defined as having a 2x fold change with a p-value < 0.05 for the comparisons between early embryos. To identify highly specific male genes, we used a 5x fold change in male over mature hermaphrodite (adult), with a p-value < 0.05 as a cutoff. Red indicates genes enriched in ES3-4, ES4-6, and males, while blue denotes genes enriched in ES1-2, ES1-3, and mature hermaphrodites. See Table S4 for the list of differentially expressed genes, their expression rpkms, and GO enrichment.

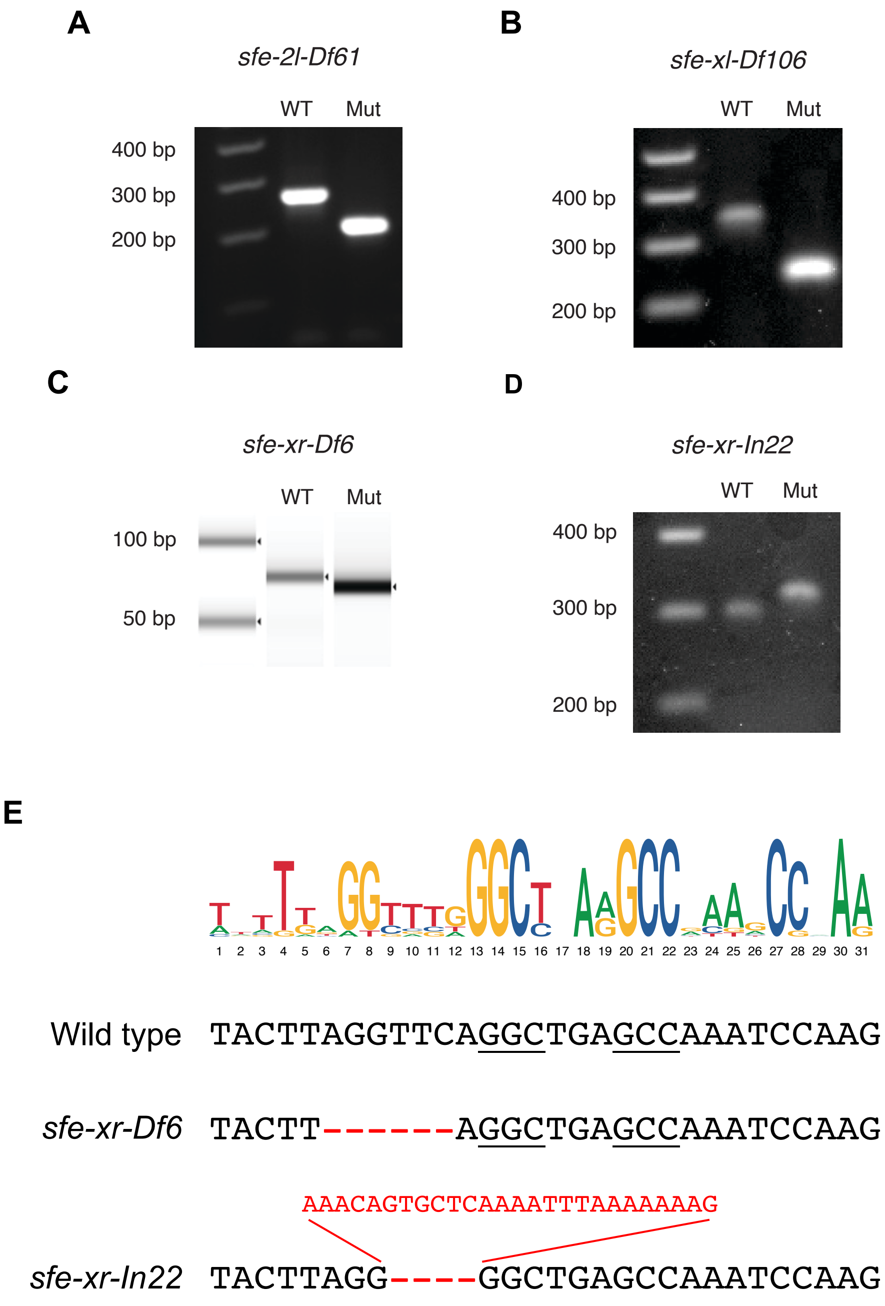


**Figure S6. CRISPR-Cas9 induced *sfe* mutations in *O. tipulae*. A-D**. PCR (A, B, D) or TapeStation (C) screenings of homozygous mutations. These mutations were confirmed with Sanger sequencing. The *sfe-xr-Df6* (C) was generated from a targeted Cas9 digestion, followed by 6-nt end resection and non-homologous end joining (NHEJ) repair. The *sfe-xr-In22* (D) was derived from a Cas9-induced break, followed by resection of 4 bp, and a defective repair process where a 26 nt segment of the repair template was used to join the chromosome ends. This resulted in 22-nt insertion. **E.** SFE motif and sequences for the wild type and the mutants (*sfe-xr-Df6* and *sfe-xr-In22*) at the right end of chromosome X. The missing or mutant bases is in red. See Table S1 for CRISPR RNAs, repair templates, and PCR primers.

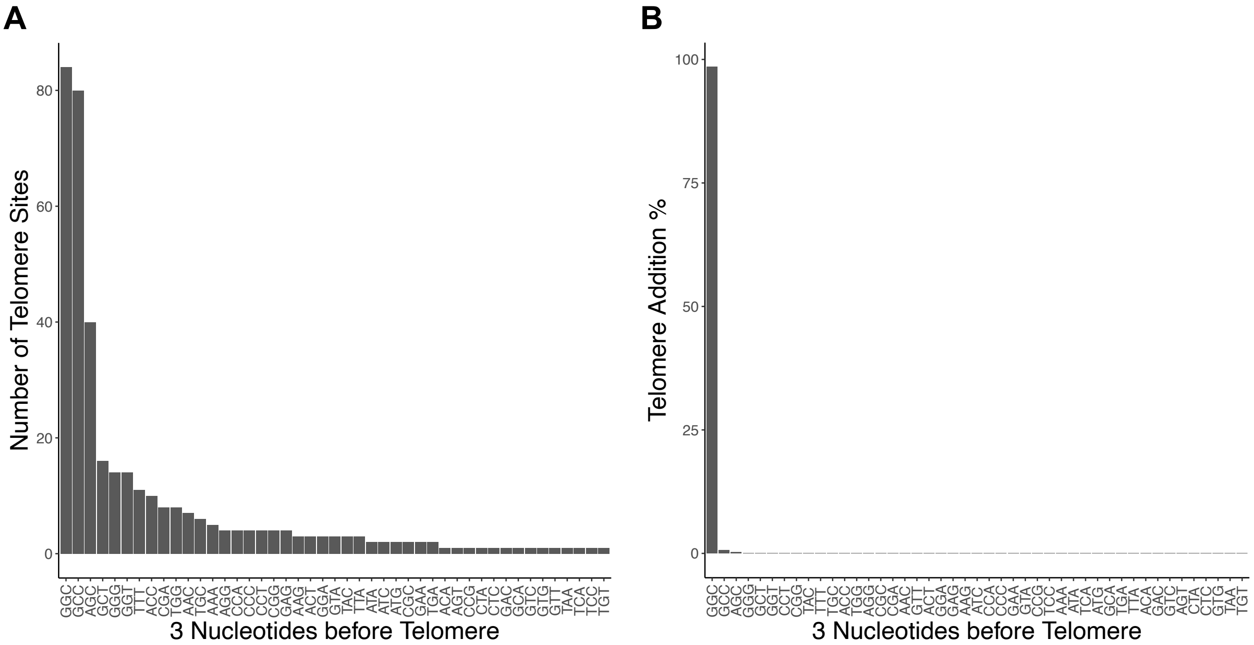


**Figure S7. Sites and frequencies for telomere addition in *O. tipulae*.** Plotted are the numbers (**A**) and frequencies (**B**) for all *de novo* telomere addition sites captured by END-seq at the 24 SFEs (Fig. 5B). The DNA for the eliminated side of the motif was reverse complemented to make all 3-mer sequences the same strand as the G-rich strand of the telomere (TTAGGC)n. Many sites can be used for telomere addition (**A**); however, the 3-nt matching GGC is the most frequent site used with 98% of the telomere reads (**B**). Most of the other 3-mers have two nucleotides that match the telomeric sequence, providing the necessary primer for telomere addition.

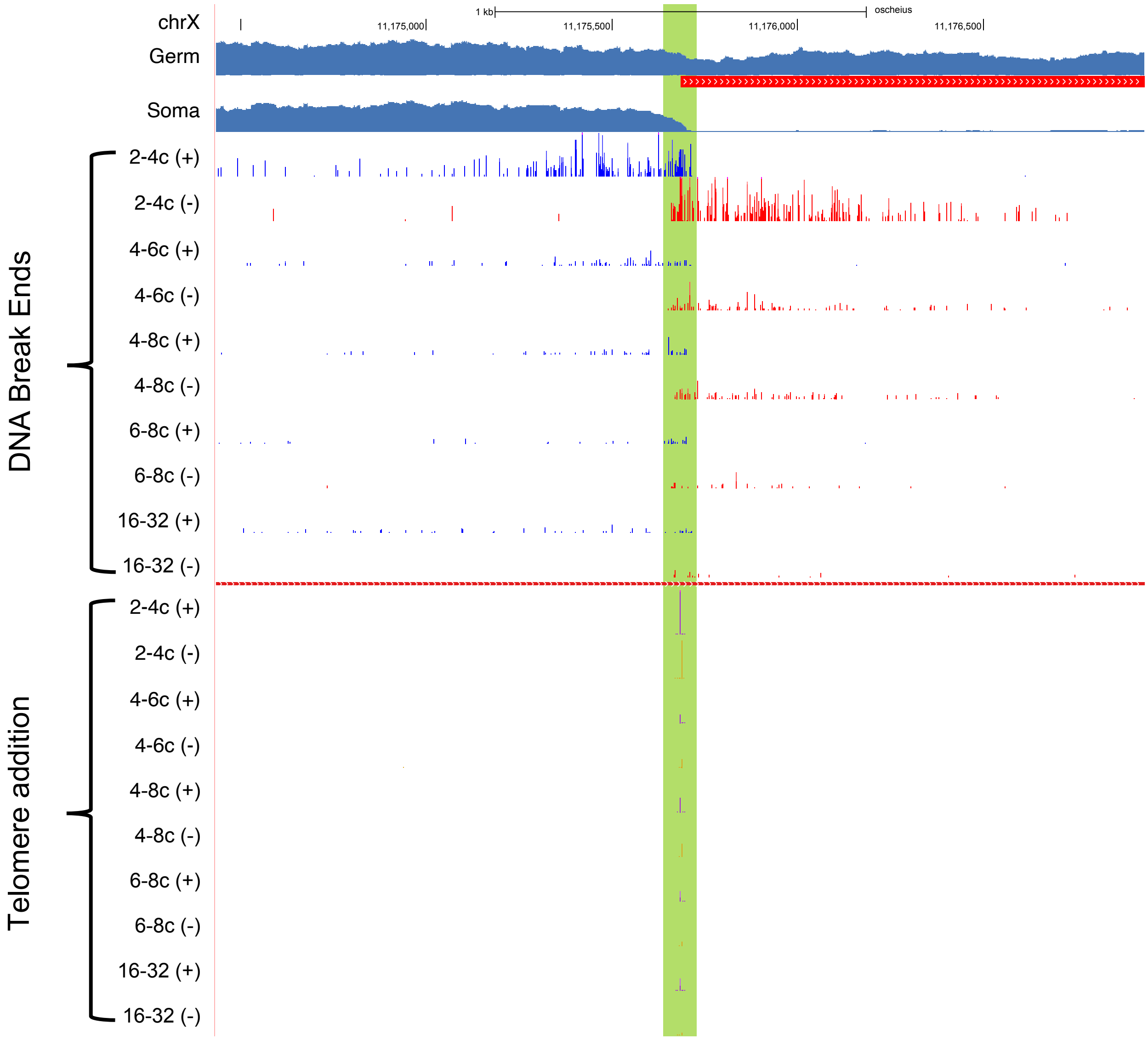


**Figure S8. DNA break and telomere addition through *O. tipulae* early embryogenesis.** A genome browser view of the right end of chromosome X, where an SFE is highlighted in green. Genomic reads coverage from germline and somatic are shown, with eliminated DNA marked in red. The top section shows the frequency of the 5'-ends of the END-seq reads, marking the blunted DSB ends. The bottom section illustrates the frequency of the telomere addition sites across the developmental stages. The data for each developmental stage is normalized and split into the forward (+) and reverse (-) strands (see Methods). Most DSBs and telomere healing is detected at the 2-4 cell stages, consistent with our prediction that PDE likely starts at the 2-cell stage (see Fig. S2).

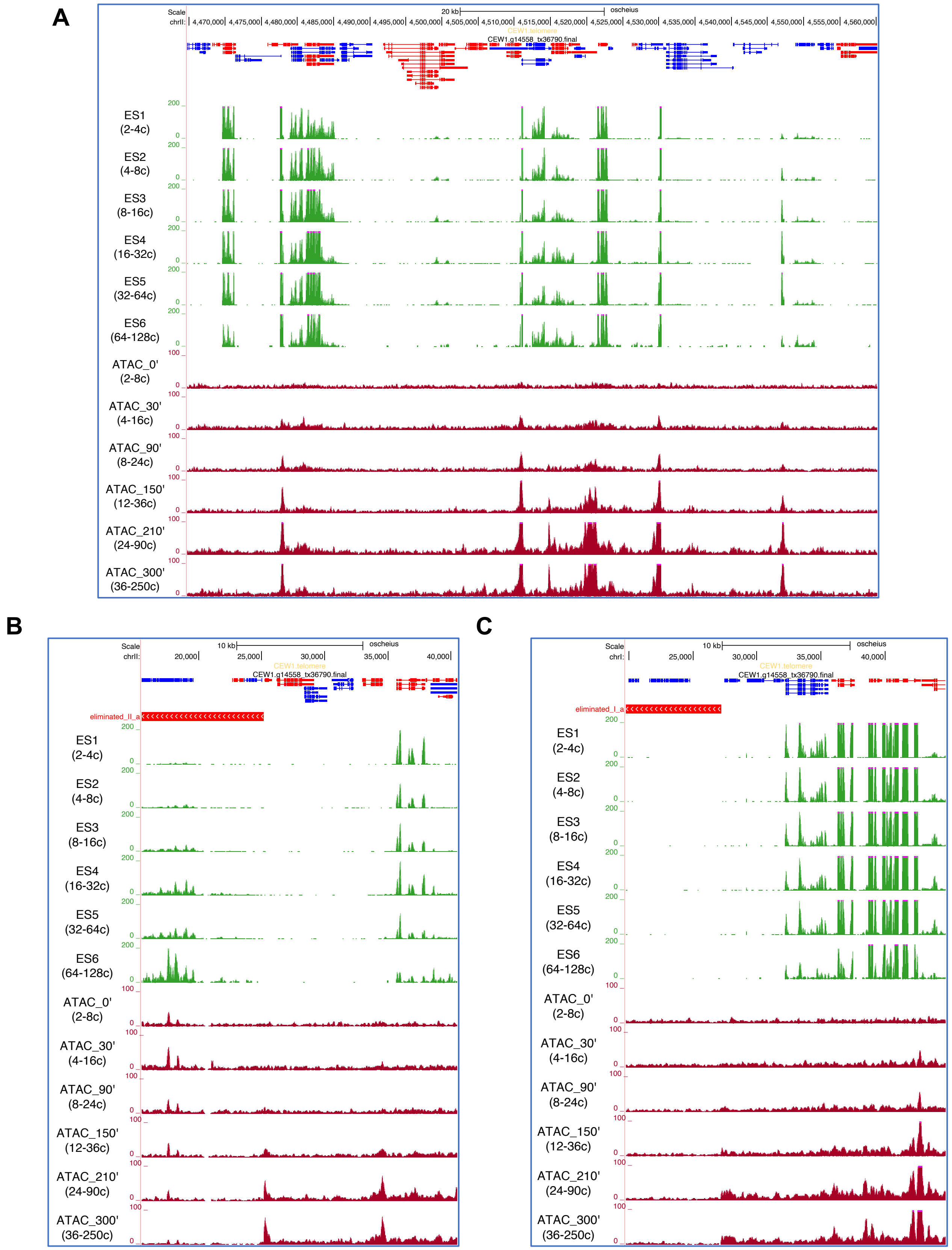


**Figure S9. ATAC-seq through *O. tipulae* early embryogenesis.** Examples of ATAC-seq (red) and RNA-seq (green) in the middle (**A**, chromosome II) and ends (**B**, chromosome II left, and **C**, chromosome I left) of the genome. No consistent patterns of chromatin accessibility were observed for the break sites or eliminated DNA.

**A** ATGAGCGACTCCATCACTTTCACCACCCGCCTCGAGGGTAAATGTTTCCAATCAGACAAAAATCGCCCCCGGGCGCACGGATCGGCATTCTCTTCTCGTA
TCTCGTCCTCTTCACTATTTCAGTTTTCCGCGCCCGTCTTTGTTCTCAAGGCATTTTCGATTCGATTTCTCCCCGACTCCGCTTTAAAAGCCGGTGGGCT
GCTGTCTCATCCTCATTTAATCACTGCTCCTTTTTCTTTCATTATCCTCATTTCCTCTGGTTACACTGCAACTTTTTAGGA**ATG**AGCGTCCAAGGGGCCA
CCTCGGGTGCCCTTCTCTTCTCCGGAGTCACTCTTCTCGTCTCCCTTTTTGCCTATGCTACCATCTACAGTCAGGTCAATGCTATCTGGGAGGAACTGGA
CCAGGAGATGGACAGCTTCAAGTCTCTGACCAACGATCTGTGGGGAGATATGGTCAAGCTGGGAGCTGGATCTGCCTCAAACCGTGTCCGT**CGT**CAGTAT
GGAGGATACGGTGCTAGTGGAGTTCAGCCTCAGCCTCCTCAGGCTCCCAACGACATCCCGGAGCAGCCTGGAAAGGACCCCGTGGGGTAAGTCGTTATA
TTTTTAAGGACTTGTGAGAGTCAGCTTGTTAAGCTGTCTGGCTCGTTACTTATTTTTTTGAGACACTTTCGTAGTCTTGTCTTTCAGCCCAACTCCAACC
CCGCTCCCTTCCCCGGATTCAATAACCCCGTTCCTCCTAGAGATCGTTGCCAGTGCACCACCGAGCACAGCTGCCCTGCCGGACCCGCTGGAGAGGCCGG
AGAGCAAGGACCCGATGGTCTTGATGGCCTTGACGGAGTAGATGGAAAAGACGGTCAGAATGCCGAAGACGTCCAGAACGAGGGAACTCACGGCTGCTTC
ACCTGCCCTCAAGGTCCTCCTGGACCTGGAGGACCTTCTGGATCCCCCGGTATGCGCGGAATGCGTGGAGCTCGAGGACAGCCCGGCCACCCTGGTCGCG
ATGGAAACCCCGGGATGCCCGGAGAGATGGGACCTCCAGGACCCCCAGGTGCCGACGGACATGTTGGAGAGCCTGGAACCAAGGGAGACGACGCTGAGCG
ACCCGTAGGTCGTGCTGGATCTCGTGGACCTCCCGGAGAGAGCGGACCCGAAGGACCCCAAGGAACTCCAGGTCGCGATGCTTACCCCGGACAGAGTGGA
CCTGTCGGAGAGCCAGGACAGCAGGGATACCAAGGAGCTGCTGGACCGGATGGAGAGGAAGGACCTCAGGGCCCTCCTGGAGCTCCTGGAAAAGACGCCA
CCTACTGCCAGTGTCCTCCAAGAGAGGAGGCTCACGGTGCCTGGAGATCTCGCAGAAAGCACCGCAAGCTCTGA

**B**

```
O. tipulae rol-6        1 MSVQGATSGALLFSGVTLLVSLFAYATIYSQVNAIWEELDQEMDSFKSLTNDLWGDMVKL 60
                          MS+ GATSGALLFSG TL+VSLFA A+IYSQV +IW ELD EMD+FKSLTNDLW DMVKL
A. rhodensis rol-6      1 MSLSGATSGALLFSGATLIVSLFAVASIYSQVTSIWNELDNEMDNFKSLTNDLWTDMVKL 60

O. tipulae rol-6        61 GAGSASNRVRRQ-YGGYGASGVQPQPPQAP 89
                           GAGSASNRVRRQ YGGYGASGV  QPP AP
A. rhodensis rol-6      61 GAGSASNRVRRQNYGGYGASGV--QPPAAP 88
```

**C**

**WT**: GAGATATGGTCAAGCTGGGAGCTGGATCTGCCTCAAACCGTGTC**CGTCGT**CAGTATGGAGGATACGGTGCTAGTGGAGTTCAGCCTCAGCCTCCTC

**rol-6**: GAGATATGGTCAAGCTGGGAGCTGGATCTGCCTCAAACCGTGTC**AGATGC**CAGTATGGAGGATACGGTGCTAGTGGAGTTCAGCCTCAGCCTCCTC

**D**

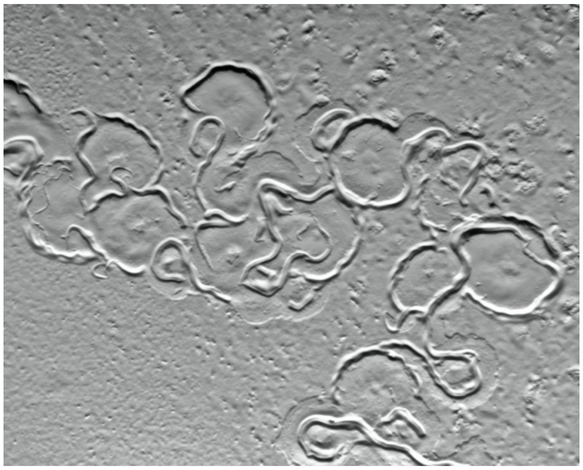

**E**

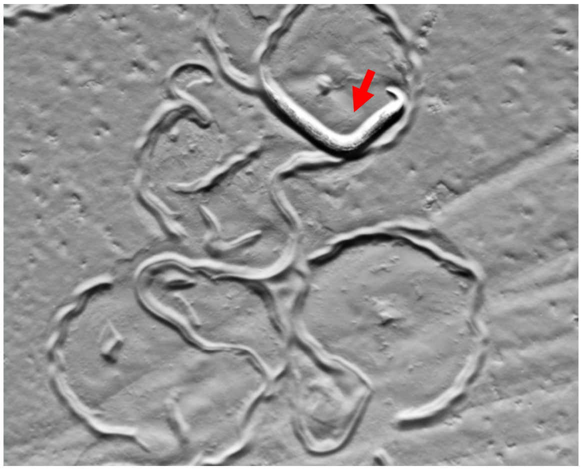

**Figure S10.** ***O. tipulae rol-6*** **gene, CRISPR, and its mutant phenotype. A.** The nucleotide sequence of *O. tipulae rol-6* mRNA (gene id: Oti_g09018). The start codon is in bold and highlighted in green. The codon specifying an arginine residue, in bold and highlighted in yellow, is mutated to cysteine to elicit a "roller" phenotype. Our RNA-seq data indicates the rol-6 gene is highly expressed in the L2 - L4 larvae stages. **B**. Protein alignment (BLASTP) of *O. tipulae* and *A. rhodensis* rol-6 protein N-terminal domains, with the to-be-mutated arginine highlighted in yellow. **C.** Wild type and the mutant rol-6 sequence repair template encoding the to-be-mutated arginine. The CGT -> AGA (in gray) is a synonymous change that prevents the crRNA (underline) from recognizing the repaired DNA. The CGT -> TGC (in yellow) is the Arg (R) to Cys (C) substitution that leads to a roller phenotype. **D** and **E.** Roller phenotype of *rol-6* mutant. **D** shows the stereotypical pattern of tracks from the mutant on an agar plate. **E** illustrates a roller worm (red arrow) with a characteristic body shape that resembles *C. elegans* roller worms. Images were taken using a Zeiss Axiozoom.v16 microscope.
